# A focal adhesion kinase-YAP signaling axis drives drug tolerant persister cells and residual disease in lung cancer

**DOI:** 10.1101/2021.10.23.465573

**Authors:** Franziska Haderk, Celia Fernández-Méndez, Lauren Čech, Johnny Yu, Ismail M. Meraz, Victor Olivas, Dora Barbosa Rabago, D. Lucas Kerr, Carlos Gomez, David V. Allegakoen, Juan Guan, Khyati N. Shah, Kari A. Herrington, Oghenekevwe M. Gbenedio, Shigeki Nanjo, Mourad Majidi, Whitney Tamaki, Julia K. Rotow, Caroline E. McCoach, Jonathan W. Riess, J. Silvio Gutkind, Tracy T. Tang, Leonard Post, Bo Huang, Pilar Santisteban, Hani Goodarzi, Sourav Bandyopadhyay, Calvin J. Kuo, Jeroen P. Roose, Wei Wu, Collin M. Blakely, Jack A. Roth, Trever G. Bivona

**Affiliations:** Department of Medicine, University of California, San Francisco; San Francisco, CA, USA; Helen Diller Family Comprehensive Cancer Center, University of California, San Francisco; San Francisco, CA, USA; Department of Cellular and Molecular Pharmacology, University of California, San Francisco; San Francisco, CA, USA; Instituto de Investigaciones Biomédicas “Alberto Sols”, Consejo Superior de Investigaciones Científícas (CSIC) y Universidad Autónoma de Madrid (UAM), Centro de Investigación Biomédica en Red de Cáncer (CIBERONC), Instituto de Salud Carlos III (ISCIII); Madrid, Spain; Department of Thoracic and Cardiovascular Surgery, The University of Texas MD Anderson Cancer Center; Houston, TX, USA; Department of Biochemistry & Biophysics, University of California, San Francisco; San Francisco, CA, USA; Department of Urology, University of California, San Francisco; San Francisco, CA, USA; Department of Pharmaceutical Chemistry, University of California, San Francisco; San Francisco, CA 94143, USA; Department of Bioengineering and Therapeutic Sciences, University of California, San Francisco; San Francisco, CA, USA; Center for Advanced Light Microscopy, University of California, San Francisco; San Francisco, CA, USA; Department of Anatomy, University of California, San Francisco; San Francisco, California, USA; Division of Medical Oncology, Cancer Research Institute, Kanazawa University, Japan; Lowe Center for Thoracic Oncology, Dana-Farber Cancer Institute; Boston, MA, USA; University of California Davis Comprehensive Cancer Center; Sacramento, CA, USA; Moores Cancer Center, University of California, San Diego; La Jolla, CA, USA; Vivace Therapeutics, Inc., 2929 Campus Drive, Suite 150; San Mateo, CA, USA; Chan Zuckerberg Biohub; San Francisco, CA, USA; Department of Medicine, Division of Hematology, Stanford University School of Medicine; Stanford, CA, USA

## Abstract

Targeted therapy is effective in many tumor types including lung cancer, the leading cause of cancer mortality. Paradigm defining examples are targeted therapies directed against non-small cell lung cancer (NSCLC) subtypes with oncogenic alterations in EGFR, ALK and KRAS. The success of targeted therapy is limited by drug-tolerant tumor cells which withstand and adapt to treatment and comprise the residual disease state that is typical during treatment with clinical targeted therapies. Here, we integrate studies in patient-derived and immunocompetent lung cancer models and clinical specimens obtained from patients on targeted therapy to uncover a focal adhesion kinase (FAK)-YAP signaling axis that promotes residual disease during oncogenic EGFR-, ALK-, and KRAS-targeted therapies. FAK-YAP signaling inhibition combined with the primary targeted therapy suppressed residual drug-tolerant cells and enhanced tumor responses. This study unveils a FAK-YAP signaling module that promotes residual disease in lung cancer and mechanism-based therapeutic strategies to improve tumor response.

## INTRODUCTION

Lung cancer, of which non-small cell lung cancer (NSCLC) is the most common subtype, is the leading cause of cancer-related mortality worldwide^1, 2^. Comprehensive molecular profiling of NSCLC has defined genetic alterations that drive tumor growth, including somatic mutations in KRAS (32.2 %), EGFR (11.3 %), and NF1 (8.3 %) as well as chromosomal fusion events involving receptor tyrosine kinases (RTKs) such as ALK, ROS1, and NTRK^3^. The development of small molecule targeted agents against these alterations has revolutionized cancer therapy given their improved clinical efficacy and safety profile compared to conventional cytotoxic chemotherapy. Prominent examples of targeted inhibitors used as first-line treatment in NSCLC are Osimertinib and Alectinib for advanced EGFR-mutant or ALK fusion-positive cancers, respectively^4, 5^. However, responses to targeted therapies are typically incomplete and residual disease containing slow cycling drug tolerant cells remains, ultimately giving rise to the development of tumors with proliferative acquired resistance that drive disease progression to which patients eventually succumb^6^. Importantly, different molecular programs have been identified in NSCLC patient specimens profiled by single cell RNA sequencing at residual disease versus at later progression (acquired resistance)^7^. Residual disease cancer cells in NSCLC are characterized by a lineage plasticity switch where adenocarcinoma cells adopt an alveolar cell-like state associated with wound healing and repair, while cancer cells with acquired resistance show an enrichment of invasion- and immune suppression- associated states^7^.

The study of drug tolerant persister cells as a residual disease model has provided important insight into the evolutionary path of drug resistance development^8–12^. Drug-tolerant persister cells are defined as a small subpopulation of cancer cells that withstand drug treatment by transitioning into a reversible state of no-to-low proliferation and evading drug-induced apoptosis^8–11^. Recent work highlighted the transcriptional co-activator YAP as an important mediator of drug tolerance by limiting pro-apoptotic BMF expression upon targeted treatment in EGFR-mutant NSCLC^13^. YAP and its paralog TAZ are effector molecules operating downstream of the canonical Hippo signaling cascade, which consists of the core MST and LATS kinases that when active inhibit YAP by enforcing its cytoplasmic retention^14^. Beyond inhibitory Hippo signaling, YAP activity can be positively promoted by a complex interplay of other pathways including signaling via Src family kinases^15^ or Rho GTPases^16^. Upon reduced Hippo signaling or alternative positive signaling input, YAP/TAZ are activated and translocate to cell nuclei where they interact with TEAD transcription factors and regulate gene expression^14^. Previous studies by our groups and other investigators showed that YAP plays important roles in cancer pathogenesis and drug resistance^16–21^. Yet, the full role of YAP in the evolution of drug tolerance and resistance and the mechanisms by which YAP can be activated to limit therapy response in NSCLC and other cancers remain incompletely understood.

## RESULTS

### Nuclear localization and activity of YAP drives drug tolerance in NSCLC

We developed several patient-derived preclinical models of residual disease to investigate the underlying mechanisms of drug tolerance. Based on established parameters of persister cells (Fig. 1a)^8–11^, we evaluated the establishment of a drug-tolerant cancer cell population across cell line models harboring different oncogenic driver mutations (Fig. 1b-e, Supplementary Fig. 1a-h). Cells were treated with their corresponding targeted inhibitor at an 80% inhibitory concentration (IC80, Supplementary Fig. 1a-c), according to prior literature on the derivation of drug-tolerant persister cells under high-dose drug treatment^8, 9, 11^. We monitored drug responses across PC9 (EGFR^del^^19^), H1975 (EGFR^L858R/T790M^), H3122 (EML4-ALK^v1^), H2228 (EML4-ALK^v3a/b^), H358 (KRAS^G12C^), and H1838 (NF1^LOF^) cells. After an initial cytotoxic response (day 1-4) marked by decreased cell culture confluency (Fig. 1b-c, Supplementary Fig. 1d-f) and increased apoptosis (Fig. 1d-e, Supplementary Fig. 1h), a subpopulation of cancer cells remained in culture at stable confluency despite continuous drug exposure (> day 5). These cells represent low-to-non-proliferative, drug-tolerant persister cells and were detected across all cell line models studied. Of note, persister cells showed a reversible phenotype regarding their lack of drug-induced apoptosis, as demonstrated by regained treatment sensitivity after drug washout (Supplementary Fig. 1h). This reversibility suggests a non-genomic mechanism(s) underlying the drug tolerance in these models. Further, suppression of oncogene-mediated signaling was maintained throughout the establishment period of persister cells, while dynamic expression changes and upregulation of known resistance-promoting proteins such as alternative RTKs including FGFR1, ErbB2, and ErbB3 as well as the anti-apoptotic protein Bcl-xL were observed (Supplementary Fig. 2a-c).

**Fig. 1.**
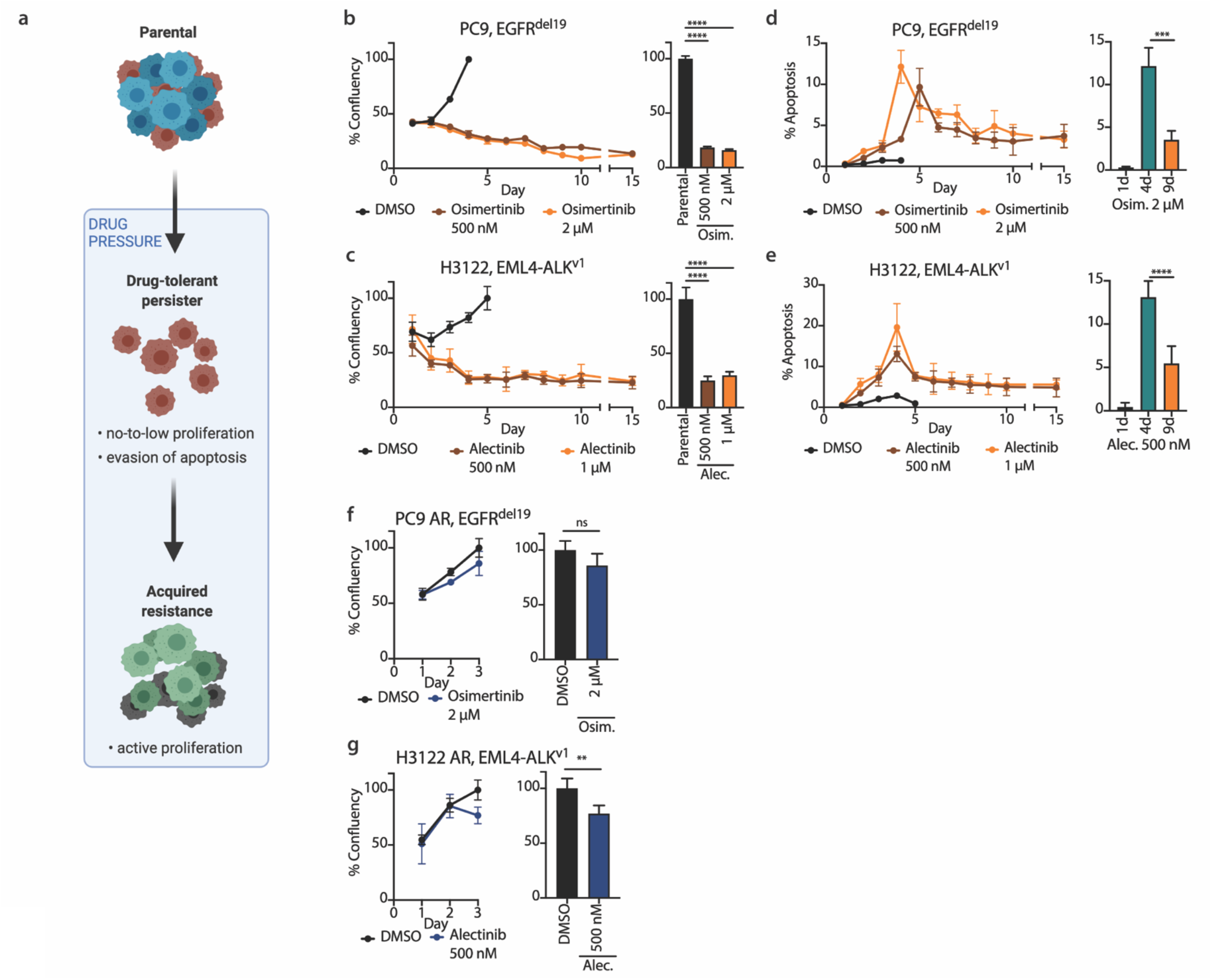
Characterization of drug-tolerant persister cells. (**a**) Schematic highlighting characteristic differences between treatment-sensitive parental cells, drug-tolerant persister cells, and acquired resistant cells. Schematic diagram was created with BioRender.com. (**b-g**) High-content microscopy screen monitoring relative cell numbers (**b-c, f-g**) and apoptosis (**d-e**) in cells treated with targeted inhibitors. (**b-c**) Confluency of EGFR-mutant PC9 cells treated with 500 nM (brown) and 2 µM (orange) osimertinib as well as of ALK-fusion positive H3122 cells treated with 500 nM (brown) and 1 µM (orange) alectinib compared to 0.1 % DMSO control (black), respectively (left), *n* = 6 per datapoint. Comparison of full cell confluency reached in 0.1% DMSO-treated parental cells to persister cell numbers at day 8 of treatment (right). Statistical evaluation by unpaired t-test. **** *p* < 0.0001. (**d-e**) Apoptosis levels in EGFR-mutant PC9 cells treated with 500 nM (brown) and 2 µM (orange) osimertinib as well as in ALK fusion-positive H3122 cells treated with 500 nM (brown) and 1 µM (orange) alectinib compared to 0.1 % DMSO control (black), respectively (left). *n* = 6 per datapoint. Comparative analysis of apoptosis levels at day 1, day 4 and day 9 of treatment (right). Statistical evaluation by unpaired t-test. *** *p* = 0.0004; **** *p* < 0.0001. (**f-g**) Confluency of osimertinib-resistant EGFR-mutant PC9-AR cells treated with 2 µM osimertinib (blue) as well as of alectinib-resistant ALK fusion-positive H3122-AR cells treated with and 1µM alectinib (blue) compared to 0.1 % DMSO control (black), respectively (left). *n* = 4 per datapoint. Comparison of cell confluency in AR cells treated with 0.1% DMSO or the respective targeted inhibitor at day 3 of treatment (right). Statistical evaluation by unpaired t-test. ns, *p* = 0.0842; ** *p* = 0.0076.

Fundamentally different from drug-tolerant persister cells, proliferation in presence of drug was regained in cells with acquired resistance that emerge upon longer-term drug exposure (> 6 weeks) (Fig. 1e-f). In addition, significant differences in gene expression were observed by RNA-seq analysis when comparing transcriptional profiles of acutely treated cells (48 h), persister cells, and cells with acquired resistance across EGFR-mutant and ALK fusion-positive cancer cell lines under therapy (Supplementary Fig. 2d-f). This highlights the concept that differential biological events can characterize each distinct treatment phase^8, 11^.

Prior work by our group and other investigators indicated a role for YAP in promoting innate and acquired resistance to targeted therapy and recent findings showed a role for YAP in drug tolerance and cancer dormancy in EGFR-mutant NSCLC^13, 20^. This prompted us to investigate the open questions as to whether YAP could function as a more general, central mediator of drug tolerance and if so, the mechanisms by which YAP is activated in this context, which are largely unknown. YAP acts as a transcriptional co-activator and can relocate to the nucleus to modulate gene expression by interacting with specific transcription factors (Fig. 2a). By performing nuclear-cytoplasmic fractionation assays, we found increased nuclear YAP in drug-tolerant persister cells across several cell line models (Fig. 2b-c). As highlighted in PC9 osimertinib persister cells, the upregulation of YAP occurs within the first 24 hours of treatment and nuclear YAP levels peak at the persister time point (Fig. 2b). Similarly, a strong increase in nuclear YAP levels was observed in H1975 osimertinib persister cells, H3122, H2228 and STE-1 alectinib persister cells as well as H358 and H1838 RMC-4550 treated (SHP2 inhibitor) persister cells (Fig. 2c). We developed isogenic, endogenously mNeonGreen-tagged YAP reporter cell lines to independently confirm the increased nuclear YAP present in drug-tolerant persister cells (Supplementary Fig. 3a). Further, given the developmental role of YAP in the control of organ size and its cell density-mediated negative regulation^22, 23^, we evaluated the increase in nuclear YAP levels in persister cells across different cell confluency (sparse, intermediate, dense) and found increased YAP nuclear localization across all cell densities during treatment (Supplementary Fig. 3b-g). Given the canonical YAP-mediated engagement of TEAD transcription factors, we expanded our analysis to evaluate the interaction between YAP and TEAD in drug-tolerant persister cells. TEAD transcription factors showed a similar nuclear enrichment in PC9 osimertinib persister cells (Supplementary Fig. 3h). A direct interaction between YAP and TEAD was confirmed by proximity ligation assays in PC9 osimertinib persisters, STE-1 alectinib persisters, and H358 RMC-4550 persisters (Fig. 2d, Supplementary Fig. 3i). Similarly, endogenous immunoprecipitation of YAP confirmed the interaction of YAP-TEAD in PC9 osimertinib persister cells (Supplementary Fig. 3j).

**Fig. 2.**
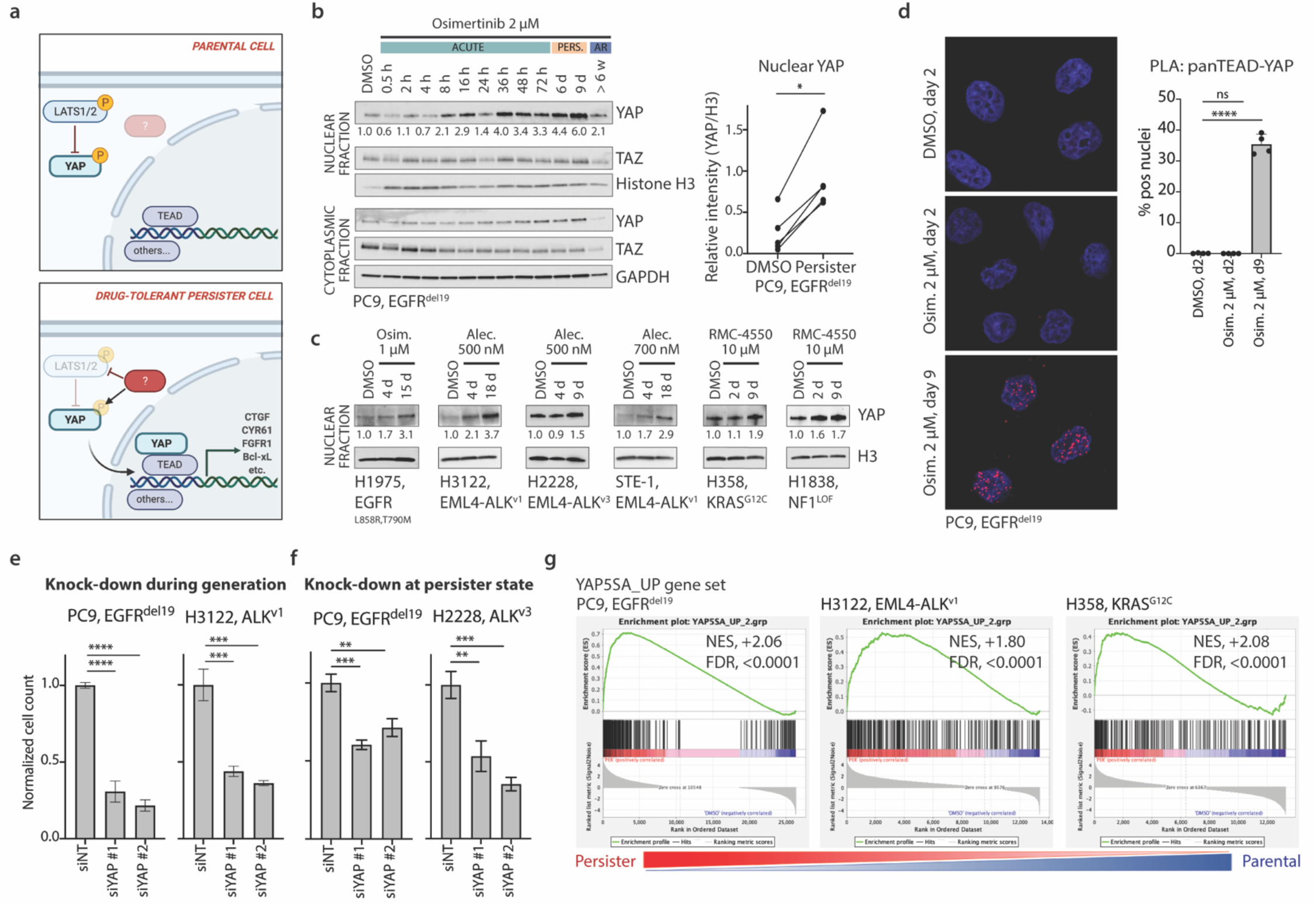
Nuclear localization of transcriptional co-activator YAP in drug-tolerant persister cells. (**a**) Schematic presentation of YAP relocalization to the cell nucleus and interaction with transcription factors in drug- tolerant persister cells. Upstream negative regulation by LATS kinase signaling is counteracted by positive signaling interactions, resulting in YAP nuclear translocation and target gene expression. Created with BioRender.com. (**b-c**) YAP levels in nuclear lysates evaluated across EGFR-mutant PC9 and H1975 cells treated with osimertinib (Osim.), ALK fusion-positive H3122 and H2228 cells treated with alectinib (Alec.) as well as KRAS-mutant H358 cells and NF1-mutant H1838 cells treated with RMC-4550. Lysates were collected from untreated cells (DMSO), acutely treated cells (day 2-4) and drug-tolerant persister cells (day 9-18). For PC9 cells, a detailed time course is presented including corresponding osimertinib-resistant PC9-AR. Quantification of nuclear YAP levels in biological replicates of PC9 persister cells compared to untreated cells is shown; *n* = 5. (**d**) PanTEAD-YAP proximity ligation assay (PLA) in EGFR-mutant PC9 cells treated with 2 µM osimertinib, analyzed by Confocal microscopy. Quantification of PLA signals per nuclei presented as mean value +/- standard deviation, *n* = 4 per condition. (**e**) Significant decrease in relative persister cell numbers upon siRNA-mediated YAP knock-down during persister cell development in PC9 cells (left) and H3122 cells (right). (**f**) Significant decrease in relative cell numbers upon siRNA-mediated YAP knock-down in PC9 osimertinib persisters (left) and H2228 alectinib persisters (right). For (**b-f**), statistical evaluation by unpaired t-test. ns, p > 0.05; *, *p* ≤ 0.05; **, *p* ≤ 0.01; ***, *p* ≤ 0.001; ****, *p* ≤ 0.0001. (**g**) Gene set enrichment analysis for the YAP-5SA_UP gene set using RNAseq expression data of EGFR-mutant PC9 cells, untreated DMSO parental control versus osimertinib persisters; of ALK fusion-positive H3122 cells, untreated DMSO parental control versus alectinib persisters; and of KRAS-mutant H358 cells, untreated DMSO parental control versus RMC-4550 persisters. *NES*, Nominal Enrichment Score; *FDR*, False Discovery Rate.

We next investigated the necessity of YAP in persister cell generation and survival. YAP silencing in parental cells suppressed the emergence of drug tolerant persister cells during therapy initiation (Fig. 2e). Moreover, YAP knock-down in persister cells caused a reduction in the number of persister cells that are maintained under drug treatment and decreased expression of survival promoting RTKs including ErbB2, ErbB3, FGFR1, and FGFR2 and the anti-apoptotic protein Bcl-xL (Fig. 2f, Supplementary Fig. 4a). Conversely, we assessed the sufficiency of YAP in limiting response to initial therapy by expressing an established mutant, hyperactive form of YAP (YAP-5SA)^24^ which shows enhanced YAP nuclear localization (Supplementary Fig. 4b). This hyperactive form of YAP was sufficient to increase drug tolerance (Supplementary Fig. 4c-h) and promote the expression of YAP responsive genes such as FGFR2 and Bcl-xL (Supplementary Fig. 4i). Expression of YAP-WT and TEAD transcription factor binding-deficient YAP-S94A showed limited or no changes across different cell line models regarding treatment response (Supplementary Fig. 4c-i). We next generated a custom gene set of transcripts upregulated in YAP-5SA-expressing PC9 cells (YAP-5SA_UP) and compared the expression profiles of these genes in parental versus persister cells. There was a significant enrichment of YAP-associated transcripts in drug-tolerant persister cells across EGFR-mutant, ALK fusion-positive, and KRAS-mutant cancer cell (Fig. 2g). Thus, YAP is hyperactivated in a conserved manner to promote gene expression changes and drug tolerance in oncogene-driven lung cancer models.

### Transcriptional adaptation enables emergence of drug persistence

Non-genetic mechanisms in the evolution of drug resistance have been highlighted in recent literature^25–28^. To address the involvement of transcriptional adaptation in YAP-driven drug tolerance, we developed isogenic EGFR-mutant NSCLC cell lines and introduced genetic barcodes of intermediate complexity (totaling 725 lineages). We conducted a single-cell RNA (scRNA) sequencing trajectory experiment that allowed us to follow cell adaptation under osimertinib treatment (Fig. 3a). Cell cycle arrest and apoptosis induction were observed during early treatment timepoints (Supplementary Fig. 5a-b), indicating drug sensitivity. Transcriptional states shifted when expression patterns were resolved over time, showing a bottleneck phase between the 8- and 24- hour timepoint and subsequent development of drug tolerant persister cells in a more similar transcriptional state (Fig. 3b). By resolving the YAP-5SA_UP gene set across the scRNA sequencing trajectory in osimertinib treated cells, we revealed the engagement of our YAP-responsive gene expression profile across individual transcripts and treatment timepoints (Fig. 3c). Thus, we were able to specify subsets of YAP-driven genes that are selected under drug pressure (Cluster 3, Fig. 3c-d). The latter showed a correlation in their onset of expression with prior observations regarding increased YAP nuclear localization within 24 hours after treatment initiation (Fig. 2b) and the defined transcriptional state of developing persister cells (Fig. 3b).

**Fig. 3.**
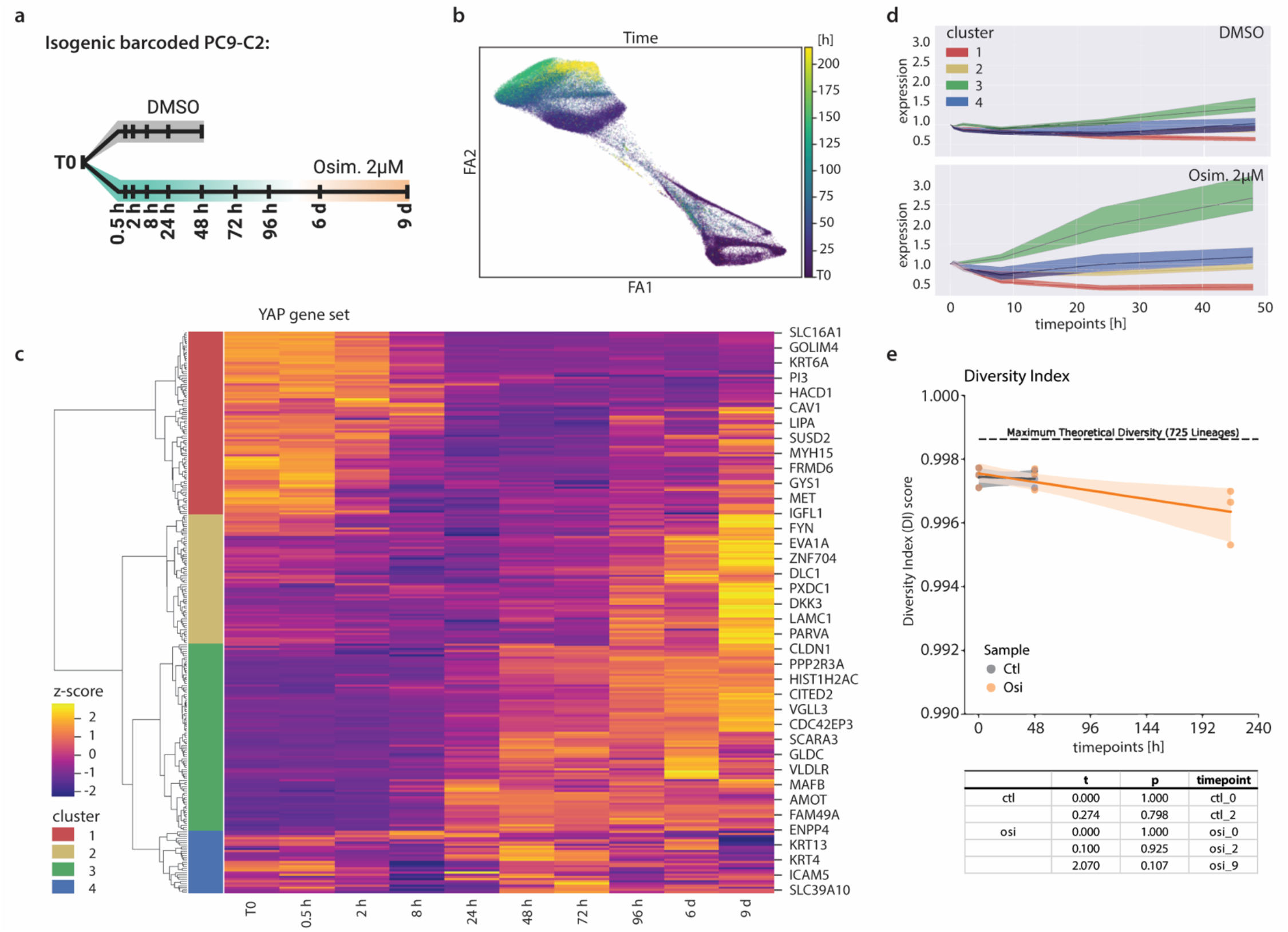
Single cell RNA sequencing trajectory in genetically barcoded, isogenic cells to identify mechanisms of transcriptional adaptation in drug tolerance. (**a**) Experimental outline indicating treatment conditions and time points for single cell RNA (scRNA) sequencing trajectory study. Schematic created with BioRender.com. (**b**) Expression changes upon treatment with osimertinib, showing the transcriptional development over the time of treatment with a noticeable shift in expression from early (t0 to 8h) to later time pointes (> 24h). (**c**) Expression changes for YAP-associated transcriptional targets (YAP-5SA_UP gene set) along trajectory in osimertinib-treated conditions, with unsupervised hierarchical clustering according to differences in expression patterns identified at single cell level. (**d**) Relative expression for YAP gene set clusters upon treatment with 0.1% DMSO control or 2 µM osimertinib, indicating selective enrichment of YAP cluster 3 gene expression under osimertinib treatment. (**e**) Diversity index based on the relative enrichment or decrease of genetic barcodes in DMSO and osimertinib trajectories with statistical evaluation demonstrating no significant selection of genetically labeled cell subsets.

By evaluating changes in the composition of genetic barcodes during treatment, no significant enrichment of cell subsets in the trajectory model was identified (Fig. 3e). This suggests that the selection of pre-existing subclones is unlikely to be a major driver of the observed phenotype of drug tolerance in this system. This was confirmed by comparing genetic shifts in PC9- and H1975-derived isogenic cell lines via whole exome sequencing, which showed identical density plots of the mutant allele frequency for cells isolated at treatment start (T0) and at persistence (9 days, Supplementary Fig. 5c-d). Further, bulk RNA sequencing of PC9- and H1975-derived isogenic cell lines validated persister cell associated phenotypes, such as a reduced cell cycle and an increased YAP-5SA_UP gene signature program (Supplementary Fig. 5e). Thus, the evolution of drug tolerance and YAP hyperactivation in these models is associated with a largely adaptive transcriptional plasticity program, within which there are specific YAP-responsive gene expression program features.

### Focal adhesion kinase signaling mediates nuclear localization of YAP during drug tolerance

In canonical Hippo pathway signaling, LATS kinases regulate YAP subcellular (cytoplasmic versus nuclear) localization such that phosphorylation of LATS on residues including Tyr1079 by upstream kinases such as MST1/2 promotes LATS kinase activity^22, 29^. Activated LATS phosphorylates YAP on serine residues such as Ser127, which results in cytoplasmic sequestration of YAP via 14-3-3 binding^22, 29^. Conversely, YAP nuclear translocation can occur upon downregulation of LATS expression or activity through various mechanisms^22, 29^. Since we observed increased nuclear YAP levels across persister cell models (Figure 2b-c), we investigated whether LATS was altered in the persister cells by measuring the levels of phospho- and total LATS. We observed decreased total LATS levels in the persister cells, while detecting modest levels and a slight increase in phospho-LATS (Tyr1079) (Supplementary Fig. 6a). Interestingly, despite downregulation of total LATS levels there was not a substantial loss of phosphorylation of YAP on Ser127 (Supplementary Fig. 6a), as might be expected upon reduction of LATS expresssion^22, 29^. These findings suggested the possibility that non-canonical regulation of YAP subcellular localization may exist in this persister cell context.

Prior work identified focal adhesion kinase (FAK) signaling as a central regulator of YAP activity in cancer cells, for which pharmacological targeting using clinical FAK inhibitors is possible^30^. Similarly, a functional relevance of FAK signaling was highlighted in resistance to the first-generation EGFR inhibitor erlotinib in EGFR-mutant NSCLC^31^. These rationales led us to investigate the hypothesis that FAK might contribute to YAP activation and drug tolerance in these oncogene-driven NSCLC systems. We first leveraged an established FAK pathway activation signature^31^ across the course of treatment in EGFR-mutant, ALK fusion-positive, and KRAS-mutant cancer cell lines. A significant increase in FAK signature gene expression during drug treatment was observed, with the highest levels in drug-tolerant persister cells (Fig. 4a). FAK gene expression changes correlated with actin remodeling upon drug treatment, resulting in increased actin polymerization and cell elongation (Fig. 4b). These findings are consistent with known roles for FAK in cytoskeletal remodeling^32^. Thus, we evaluated phosphorylation and activation of FAK and associated tyrosine kinases within the FAK signature, i.e., EphB1 and ACK1, as combinatorial knock-down of all three molecules had been reported to mediate cell death in EGFR inhibitor-resistant cells^31^. Indeed, an increase in phosphorylation for EphB1, ACK1, and FAK was observed in PC9 osimertinib persister cells and H3122 alectinib persister cells compared to parental cells (Fig. 4c). FAK was further reported to mediate YAP phosphorylation at Tyr357 and to increase YAP activity^30^. As expected, YAP- activating phosphorylation at Tyr357 and higher YAP stability were detected in drug-tolerant persister cells, while limited alterations in canonical LATS phosphorylation and inhibitory YAP Ser127 phosphorylation were observed (Fig. 4c). Blocking critical FAK-associated signaling by combinatorial knock-down of FAK, ACK1 and EphB1 in persister cells resulted in a significant reduction of persister cell viability (Fig. 4d) and was characterized by a decrease in YAP nuclear localization and overall YAP levels (Fig. 4e). Similarly, CRISPR-mediated FAK knock-out (KO) significantly suppressed the emergence of drug-tolerant persister cells, with similar effects as seen upon direct YAP-KO (Fig. 4f). Further, FAK-KO PC9 cells showed a significant reduction of YAP nuclear levels (Fig. 4g). Pharmacological inhibition of FAK signaling by combinatorial treatment with VS-4718, an established potent and selective FAK inhibitor^33^ (Fig. 4h, Supplementary Fig. 6b), or the multikinase inhibitor dasatinib, that targets SRC downstream of FAK^34^ (Supplementary Fig. 6b-c), in combination with the primary oncoprotein targeted therapy resulted in a significant reduction of persister cell viability, while limited single agent activity of VS-4718 or dasatinib in parental cells was noted (Supplementary Fig. 6d). VS-4718 treatment reduced YAP nuclear levels in osimertinib-treated PC9 cells and alectinib-treated H3122 cells (Fig. 4i) and decreased YAP Tyr357 phosphorylation (Supplementary Fig. 6b). The collective findings suggest an unanticipated, critical role for FAK signaling engagement in promoting YAP nuclear localization and persister cell evolution and survival in oncogene-driven NSCLC systems. The data indicate that FAK signaling represents a distinct targetable axis to suppress persister cell survival.

**Fig. 4.**
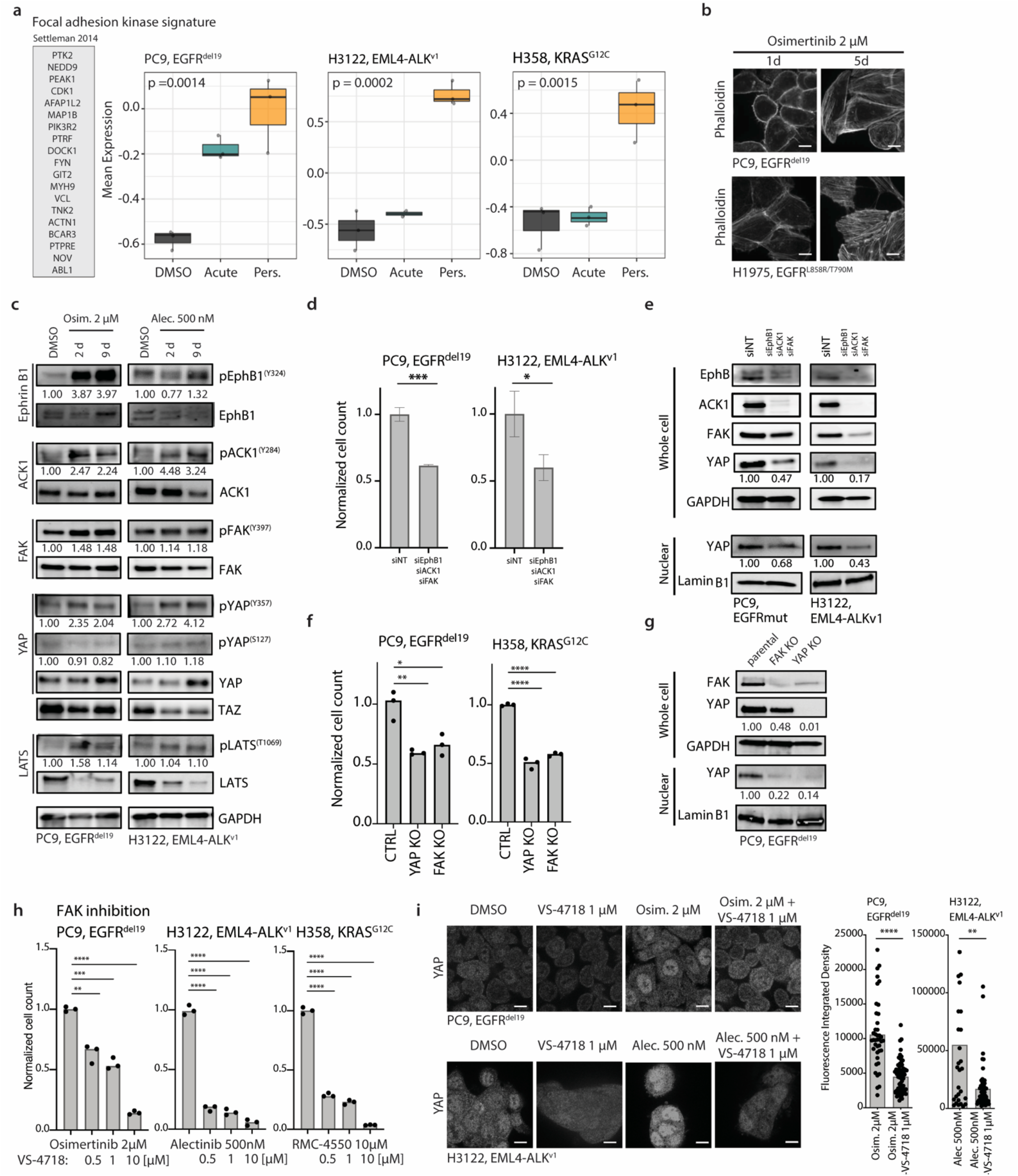
Upstream regulation of YAP nuclear enrichment by FAK signaling in drug-tolerant persisters. (**a**) Changes in the FAK expression signature^31^ for EGFR-mutant PC9 cells treated with 2 µM osimertinib, ALK fusion-positive H3122 cells treated with 500 nM alectinib, and KRAS-mutant H358 cells treated with 10 µM RMC-4550. Statistical significance is indicated by two-way Anova test. (**b**) Actin cytoskeleton changes upon treatment with 2 µM osimertinib in EGFR-mutant PC9 and H1975 cells, scale bar: 10 µm. (**c**) Phosphorylation changes of key FAK signaling molecules including EphB1, ACK1, and FAK, as well as for the YAP activating Y357 and inactivating S127 phosphorylation site upon treatment with targeted inhibitors in EGFR-mutant PC9 cells and ALK fusion-positive H3122 cells. (**d**) Relative persister cell number upon combinatorial knock-down of EphB1, ACK1, and FAK (siEAF) in PC9 osimertinib persister cells and H3122 alectinib persister cells compared to non-target control (siNT). (**e**) Changes in YAP expression and nuclear localization upon combinatorial knock-down siEAF in PC9 osimertinib persister cells and H3122 alectinib persister cells. (**f**) Relative persister cell number in EGFR-mutant PC9 cells and KRAS-mutant H358 cells harboring a CRISPR- mediated FAK or YAP knock-out (KO). EGFR-mutant PC9 cell lines have been treated with 2 µM osimertinib. KRAS-mutant H358 cell lines have been treated with 10 µM RMC-4550. (**g**) Changes in YAP expression and nuclear localization upon FAK KO in parental EGFR-mutant PC9 cells. (**h**) Normalized persister cell numbers upon targeted therapies in combination with FAK inhibitor VS-4718 across EGFR-mutant PC9 cells treated with 2 µM osimertinib, ALK fusion-positive H3122 cells treated with 500 nM alectinib, and KRAS-mutant H358 cells treated with 10 µM RMC-4550. (**i**) Changes in YAP nuclear localization in EGFR-mutant PC9 cells (day 5) and ALK fusion-positive H3122 cells (day 2) upon combination treatment with FAK inhibitor VS-4718 and the backbone targeted treatment; scale bar: 10 µm. Quantification of relative integrated density was performed by automated analysis quantifying the intensity for the protein of interest per nuclei. For (**d**, **f**, **h-i**), statistical evaluation by unpaired t-test. ns, p > 0.05; *, *p* ≤ 0.05; **, *p* ≤ 0.01; ***, *p* ≤ 0.001; ****, *p* ≤ 0.0001.

### Treatment studies in patient-derived organoid and xenograft models confirm YAP engagement in residual tumor cells

Patient-derived organoid models have been shown to recapitulate determinants of treatment response of patient specimens^35–37^, with recent reports focusing on the development of 3D NSCLC organoid cultures^38, 39^. We established two EGFR-mutant NSCLC organoid cultures from clinical patient specimens and confirmed the presence of the oncogenic EGFR driver mutation (Supplementary Fig. 7a-b). Both EGFR-mutant NSCLC organoid cultures, showed sensitivity to treatment with 100 nM osimertinib (Supplementary Fig. 7c-d) and thus, were determined therapy-responsive and suitable for persister cell generation. We established persister cell derivatives in these organoid cultures (Fig. 5a) and observed an increase in the expression of YAP transcriptional targets in the derived persister cells. (Fig. 5b, Supplementary Fig. 7e). In addition, sensitivity to combinatorial treatment with the FAK inhibitor VS-4718 along with the primary oncoprotein targeted therapy was demonstrated in EGFR-mutant NSCLC organoid culture TH107 (EGFR^del^^19^, Fig. 5c), extending our findings described above in the other systems.

**Fig. 5.**
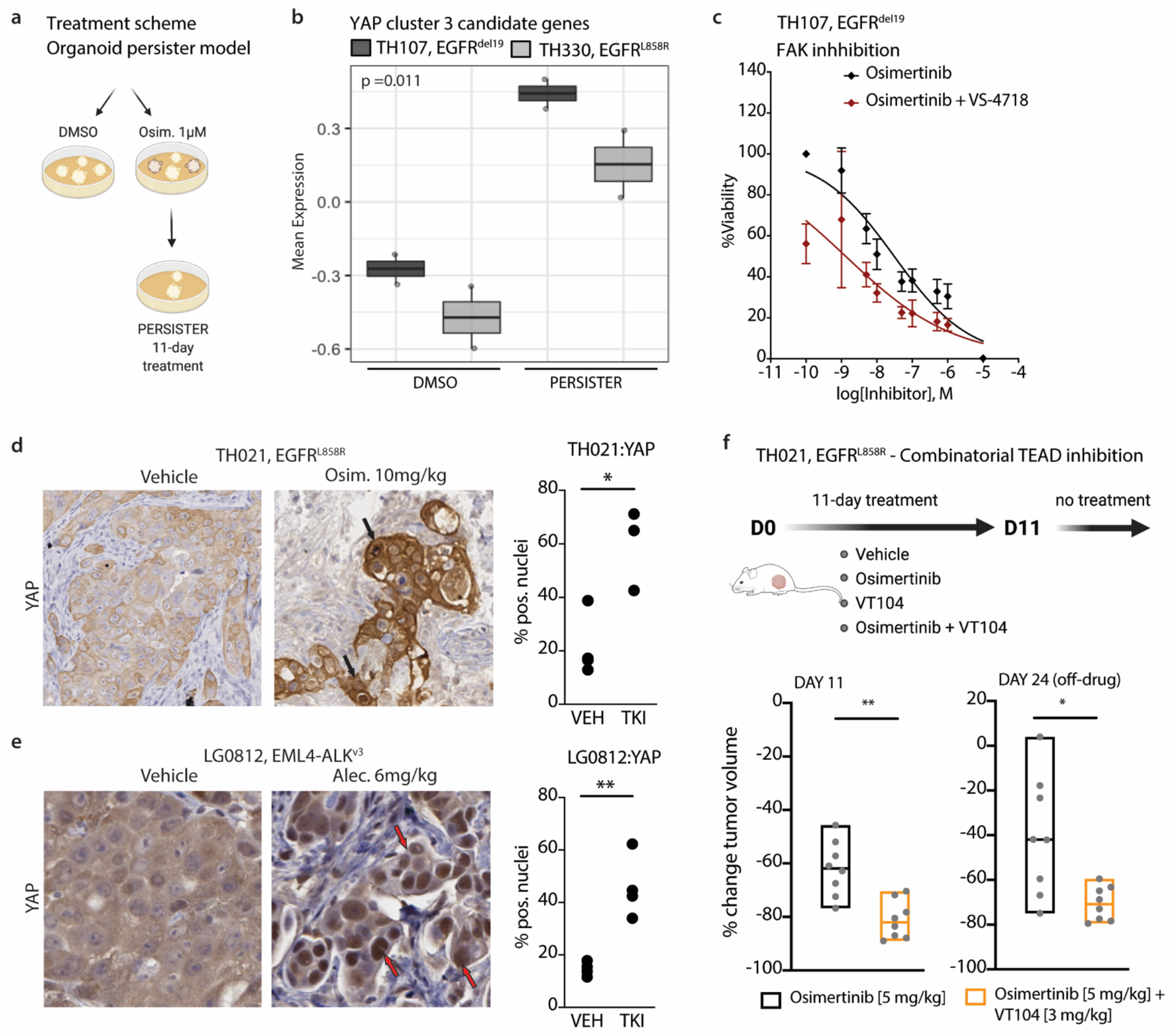
Characterization of residual disease in patient-proximate model systems. (**a**) Schematic presentation for the generation of persister cells in NSCLC patient-derived organoid (PDO) models. (**b**) Mean expression of YAP-5SA cluster 3 genes in 0.1% DMSO control (DMSO) and drug-tolerant persisters (PERSISTER) across treatment-sensitive EGFR-mutant PDO models, i.e., EGFR^del^^19^ TH107 and EGFR^L858R^ TH330. Statistical significance is indicated by two-way Anova test. (**c**) Treatment response to escalating doses of osimertinib upon combinatorial treatment with FAK inhibitor VS-4718 (1 µM) in EGFR^del^^19^ TH107 as determined by CellTiter-Glo assay. (**d-e**) Immunohistochemistry staining for YAP in residual tumors cells of EGFR-mutant TH021 and ALK fusion-positive LG0812 PDX models upon treatment with targeted inhibitors (TKI) compared to vehicle control (VEH). Quantification of nuclear levels (% nuclear) by automated image analysis. Statistical evaluation by unpaired t-test. For TH021: VEH vs TKI, * *p* = 0.0127. For LG0812: VEH vs TKI, ** *p* = 0.0022. Arrows indicate cluster of YAP positive tumor cell nuclei. (**f**) Combinatorial osimertinib [5 mg/kg] plus TEAD inhibitor VT104 [3 mg/kg] treatment in the EGFR-mutant TH021 PDX model. Treatment schematic and relative changes in tumor volumes at day 11 (on-drug) and at day 24 (off-drug) are presented. Statistical evaluation by Mann- Whitney test. ** *p* = 0.0019, * *p* = 0.0148. Schematic diagram was created with BioRender.com.

We expanded this analysis using patient-proximate *in vivo* models by first measuring YAP in treatment-sensitive NSCLC PDX samples. We treated EGFR-mutant PDX TH021 (EGFR^L858R^) with 10 mg/kg osimertinib for 7 days and ALK fusion-positive PDX LG0812 (EML4-ALK^v^^3^) with 6 mg/kg alectinib for 17 days. Both models showed sensitivity to targeted inhibition resulting in significant regression of tumor volumes under treatment (Supplementary Fig. S7f-g). However, residual tumor lesions remained at treatment endpoint. Immunohistochemistry (IHC) staining of residual tumor specimens demonstrated increased YAP nuclear levels in residual tumor cells present at the treatment endpoint in both PDX models (Fig. 5d-e). Further, RNA sequencing demonstrated a significant induction of YAP-mediated transcriptional changes in inhibitor-treated residual tumors (Supplementary Fig. 7h-i), confirming the relevance of prior findings in *in vivo* settings. In addition, combinatorial treatment with the FAK inhibitor VS-4718 along with the primary oncoprotein targeted therapy showed increased therapeutic efficacy in the EGFR-mutant H1975 (EGFR^L858R/T90M^) xenograft model that showed evidence of increased nuclear YAP levels upon EGFR inhibitor treatment alone (Supplementary Fig. 8).

In addition to inhibiting FAK signaling upstream of YAP, we also investigated therapeutic options to target the interaction of YAP with its transcription factor TEAD. As outlined previously, canonical YAP-TEAD engagement was significantly enriched in drug-tolerant persister cell models (Fig 2d, Supplementary Fig. 3h-j). Using the novel *in vivo*-ready TEAD inhibitor VT104^40^, short-term combinatorial treatment along with the primary oncoprotein targeted therapy resulted in the reduced expression of YAP signature genes in PC9 osimertinib persister cells and H3122 alectinib persister cells (Supplementary Fig. 9a). Persister cell survival was substantially impaired upon combinatorial treatment with VT104 plus the backbone oncoprotein targeted therapy, while no single agent activity for VT104 was noted (Supplementary Fig. 9b-c). Similarly, combinatorial treatment with the TEAD inhibitor VT104^40^ showed a significantly heightened tumor response at residual disease and reduced regrowth of tumors after drug withdrawal in the EGFR-mutant PDX model TH021 (Fig. 5f, Supplementary Fig. 10a). In addition, an increased therapeutic efficacy of combinatorial treatment with the TEAD inhibitor VT104 was observed in the EGFR-mutant H1975 (EGFR^L858R/T90M^) xenograft model (Supplementary Fig. 8a) in which we previously observed YAP engagement in response to osimertinib monotherapy (Supplementary Fig. 8b), without evidence of substantial overall toxicity in initial studies (Supplementary Fig. 8c).

### Humanized murine models confirm YAP-mediated drug tolerance and highlight a role for YAP in modulating treatment-mediated changes in the humanized tumor microenvironment

Prior work highlighted the complex interplay between YAP and the tumor microenvironment (TME), where YAP may both foster tumor cell survival and promote immunity against tumor cells^41^. Thus, we expanded our work to humanized murine models derived from fresh cord blood CD34+ stem cells and showing a functional immune cell repertoire in the presence of lung tumor xenograft and PDX models^42^. Implantation of EGFR-mutant PC9 parental cells, YAP-WT, and YAP-5SA overexpressing PC9 cells into humanized mice was performed to establish tumors. We monitored response to osimertinib treatment and found increased drug tolerance was mediated by overexpression of YAP-WT or YAP-5SA (Fig. 6a-b, Supplementary Fig. 10b). This confirms previous *in vitro* results in YAP cell line models (Supplementary Fig. 4c-h) and highlights a similar drug tolerance phenotype in the presence of a functional immune microenvironment *in vivo*. Of note, important changes in the cellular composition of the tumor microenvironment were observed upon osimertinib treatment. In mice implanted with PC9 parental cells, a treatment-mediated increase in tumor-infiltrating myeloid cells and T lymphocytes was observed at treatment endpoint (Fig. 6c-i). Macrophage populations were skewed towards elevated numbers of pro-inflammatory HLA-DR+/CD163- M1 type macrophages upon osimertinib treatment in the PC9 parental tumor cohort (Fig. 6c-d, Supplementary Fig. 9c). By comparison, M1 type macrophages were reduced in vehicle and osimertinib-treated groups across YAP-WT and YAP-5SA overexpressing cohorts, with the most pronounced phenotype for YAP-5SA overexpressing PC9 cells (Fig. 6c-d, Supplementary Fig. 9c). On the other hand, the abundance of tumor-supportive HLA-DR-/CD163+ M2 type macrophages were increased in untreated groups of YAP overexpressing cohorts compared to PC9 parental tumors (Fig. 6e, Supplementary Fig. 9c). However, osimertinib treatment resulted in a reduction of M2 macrophages and similar abundance across parental and YAP overexpressing groups upon treatment (Fig. 6e, Supplementary Fig. 9c). In addition to changes in the composition of myeloid cell infiltrates, complex alterations in the abundance and phenotype of infiltrating T lymphocytes were observed (Fig. 6f-i). While an increase of infiltrating T lymphocytes was detected upon osimertinib treatment in PC9 parental tumors, lower levels of infiltrating T lymphocytes (YAP-5SA, Fig. 6g) or redistribution of CD4:CD8 T cell ratios in favor of non-cytotoxic CD4+ T cells (YAP-WT, Fig. 6h-I, Supplementary Fig. 9c) were observed upon treatment in YAP overexpressing cohorts. In conclusion, YAP upregulation resulted in a shift towards a more tumor-supportive tumor microenvironment, with reduced levels of pro-inflammatory M1 type macrophages and cytotoxic T cells.

**Fig. 6.**
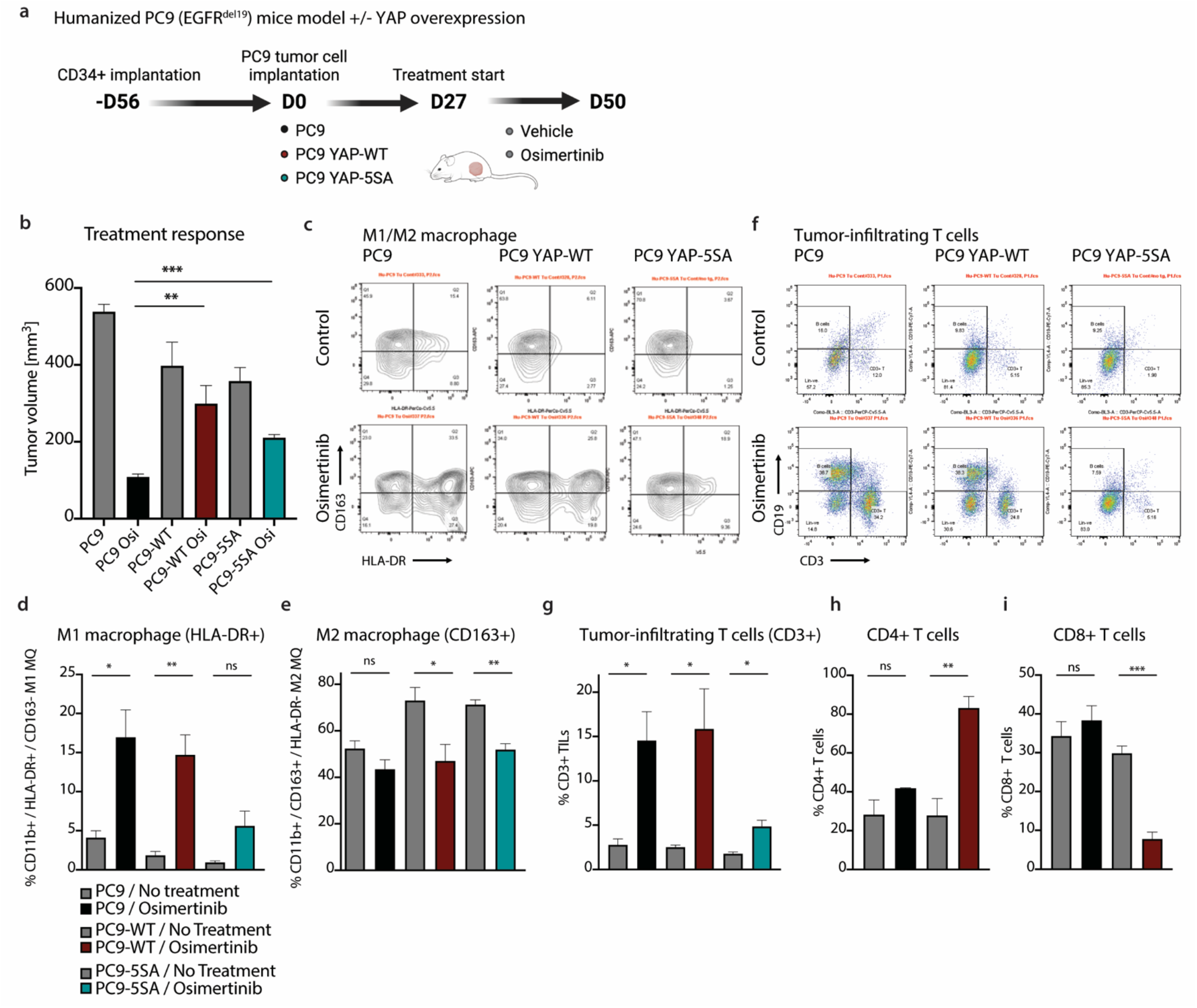
Evaluation of YAP-mediated drug tolerance in immune-competent humanized mice. (**a**) Schematic presentation for the establishment and the treatment study of the humanized EGFR-mutant PC9 xenograft model. Schematic diagram was created with BioRender.com. (**b**) Changes in tumor volume at treatment endpoint. A Osimertinib [5 mg/kg] treatment study in the humanized PC9 mouse model was conducted comparing parental cells, cells expressing YAP-WT and cells expressing hyperactive YAP-5SA. (**c**-**e**) Changes in tumor infiltrating macrophage populations at treatment endpoint. Macrophage populations are defined as CD11b+ cells, with HLA- DR+ for M1 macrophages and CD163+ for M2 macrophages. (**f**-**i**) Changes in tumor infiltrating T cell populations at treatment endpoint. T-cell populations are defined as CD25+/CD3+ cells, with differentiation of CD4+ and cytotoxic CD8+ T-cells. For **b**-**i**, statistical evaluation by unpaired t-test. * *p* ≤ 0.05, ** *p* ≤ 0.01, *** *p* ≤ 0.001.

### Single-cell RNA sequencing (scRNAseq) of NSCLC patient specimens indicates a YAP transcriptional signature at residual disease and potential therapeutic vulnerabilities

We next investigated the clinical relevance of our preclinical findings highlighting a FAK-YAP signaling axis as a driver of residual disease in NSCLC. We leveraged clinical specimens of NSCLC samples that were obtained from patients before initiating systemic targeted therapy (TKI naive [TN]), at the residual disease (RD) state, which includes samples taken at any time during treatment with targeted therapy while the tumor was regressing or stable by clinical imaging [RD], and upon subsequent progressive disease as determined by clinical imaging, at which point the tumors showed acquired drug resistance (progression [PD]). Previous work from our group highlighted the relevance of scRNAseq profiling to identify clinical treatment state-specific transcriptional programs present in cancer cells in these NSCLC samples obtained from patients collected at the 3 different treatment states^7^. For example, cancer cells remaining at residual disease specifically manifested an alveolar type 1 and 2 cell-like state associated with a more indolent state as a prelude to the subsequent acquisition of full drug resistance and re-establishment of active proliferation^7^. We used this scRNAseq expression dataset to assess the clinical relevance of YAP transcriptional activity at the residual disease time point, particularly focusing our analysis on NSCLC specimens obtained from the lungs of advanced-stage patients to avoid potential organ-site specific confounding effects. Transcriptional signatures differed across treatment response states (Supplementary Fig. 11a), with a significant increase in expression of a subset of YAP transcriptional targets in cancer cells present at residual disease compared to treatment naïve and progressive disease timepoints (Fig. 7a). Similarly, differential transcriptional programs induced upon overexpression of hyperactive YAP-5SA as well as in EGFR- mutant PC9 and ALK-fusion positive H3122 persister cells showed predominant expression features at residual disease in patients, as determined by permutation analysis (Supplementary Fig. 11b). YAP-associated transcriptional targets that are differentially expressed at residual disease in patients include the Cbp/p300 transactivator CITED2, Axl receptor agonist GAS6, TGFβ receptor ligand BMP4, AT1/AT2 alveolar marker AGER as well as proteins associated with cytoskeletal signaling, such as DLC1 as well as TNNC1 (Fig. 7b, Supplementary Fig. 11c). Similarly, assessment of YAP levels by IHC confirmed a significant increase of nuclear YAP in tumor cells present in tumor specimens at residual disease compared to TKI-treatment naïve samples (Fig. 7c, Supplementary Fig. 11d). To evaluate whether the YAP transcriptional signature is a clinically relevant biomarker of patient survival, we tested the association of YAP signature expression status and overall survival in The Cancer Genome Atlas (TCGA) lung adenocarcinoma bulk RNA-seq dataset, as previously for the AT1/2- like transcriptional signature^7^. This clinical cohort, while not comprised of many TKI-treated samples, allows for long-term follow up and survival analysis and can inform on the biological implications of genetic programs more broadly in lung cancer. We evaluated the association of YAP-mediated gene expression with disease progression and found a less aggressive phenotype in the TCGA tumor specimens showing a YAP-high expression profile (Fig. 7d, Supplementary Fig. 11e). This is similar to our observations for the AT1/AT2-like signature state present in the cancer cells at residual disease presented in prior work^7^. Further, YAP-high expression samples correlated with Cell cycle-low and FAK signature-high expression states in this TCGA cohort (Fig. 7e, Supplementary Fig. 11e). The data suggests additional clinical relevance for these characteristic persister cell features of low proliferation and high FAK-YAP signaling even beyond on-treatment residual disease and a potential link with the more indolent AT1/2-like lineage state present in NSCLCs that we showed previously.

**Fig. 7.**
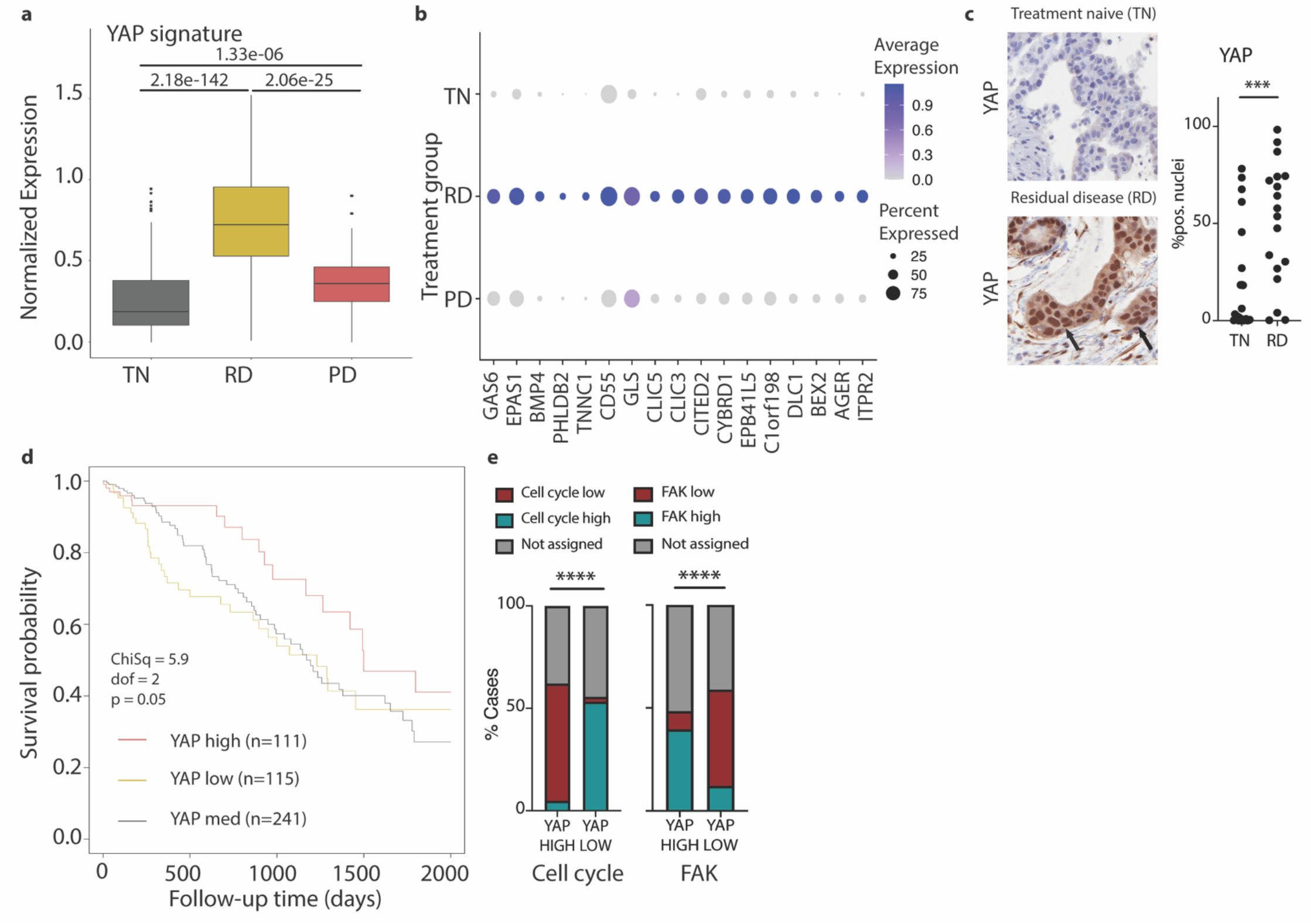
Engagement of FAK-YAP transcriptional program in patient specimen of residual disease. (**a**) Normalized expression of YAP signature genes across patient specimen classified as TN, RD or progressive disease (PD) using previously published single cell RNAseq (scRNAseq) data^7^. The YAP signature is defined by genes within the YAP-5SA_UP gene set that are significantly upregulated across PC9, H3122, and H358 persister models as well as differentially upregulated in the RD treatment timepoint. Adjusted *p*-values corrected for multiple testing by Bonferroni are presented. (**b**) Average expression of individual YAP signature genes highlighted in **A** across TN, RD, and PD treatment groups. (**c**) Immunohistochemistry staining for YAP in patient specimens classified as TKI treatment naïve (TN) or collected at residual disease upon treatment with targeted inhibitors (RD). Quantification of nuclear levels (% nuclear) by automated image analysis. Statistical evaluation by unpaired t-test. *** *p* = 0.0007. Arrows indicate cluster of YAP positive tumor cell nuclei. (**d**) Kaplan-Meier plot of the relationship between the YAP signature and patient overall survival (OS) within the TGCA-LUAD dataset. Patients were stratified by signature expression quartile (YAP low, Q1 = 155; YAP high, Q4 = 172.5; see also Supplementary Fig. 12d). Statistical evaluation by Chi-square test. (**e**) Overlap of YAP signature with Cell cycle signature (left) or FAK signature (right) in the TCGA dataset. Patients were stratified by signature expression quartile as outlined in Supplementary Fig. 12d. Statistical evaluation by Chi-square test.

Finally, we aimed to generate a systematic framework for identifying pharmacologic agents that could potentially reverse expression signatures identified in residual disease clinical specimens. This general approach was recently demonstrated in cell line models, with several inhibitors identified that correlated with a reversal of persister cell transcriptional changes and were confirmed to show sensitivity upon combinatorial treatment along with the primary oncoprotein targeted therapy^43^. We used the NIH Library of Integrated Network-Based Cellular Signatures (LINCS)^44^ L1000 transcriptomic platform as a reference dataset for perturbation-induced gene expression changes. This resource includes 71 cell lines and over 20,000 pharmacologic/chemical perturbagens. We identified 10,917 unique drug-cell line combinations with a significant correlation (positive/similar or negative/dissimilar) with NSCLC residual disease-associated gene expression changes (Fig. 8a). By selecting for drugs that were associated with opposite expression patterns compared to residual disease transcriptional profiles (negative correlation, *n* = 3,047) and limiting to pharmacological agents with target information (annotated), we identified over 1,691 drug-cell line combinations that indicate a reversal of NSCLC residual disease-associated expression patterns (Fig. 8a, Supplementary Fig. 12a-c). Pharmacologic perturbagens that induced gene expression changes that were most significant negatively correlated with the gene expression changes present in residual disease cancer cells include the JAK1/2 inhibitor momelotinib, SYK inhibitor tamatinib and SRC inhibitor dasatinib (Fig. 8a, Supplementary Fig. 12b-c). Selective FAK inhibitors within the LINCS L1000 database were limited to PF-562271^45^, which also resulted in a significant negative concordance score. Similarly, tankyrase inhibitor XAV-939 presented a negative concordance score and was predicted to reverse NSCLC residual disease-associated gene expression changes. This is interesting as XAV-939 was previously reported to suppress YAP nuclear localization and transcriptional activity^46^ as well as to increase treatment response in EGFR-mutant and ALK fusion-positive upon combination with the primary oncoprotein targeted therapy ^7^. Notably, we demonstrated that FAK inhibitor VS-4718 and SRC inhibitor dasatinib showed sensitivity in drug tolerant persister cells in this manuscript (Fig. 4h, Supplementary Fig. 6b), providing evidence of the general utility of this computational framework for identifying persister cell vulnerabilities and strengthening the relevance of FAK signaling and potentially other signaling networks in residual disease.

**Fig. 8.**
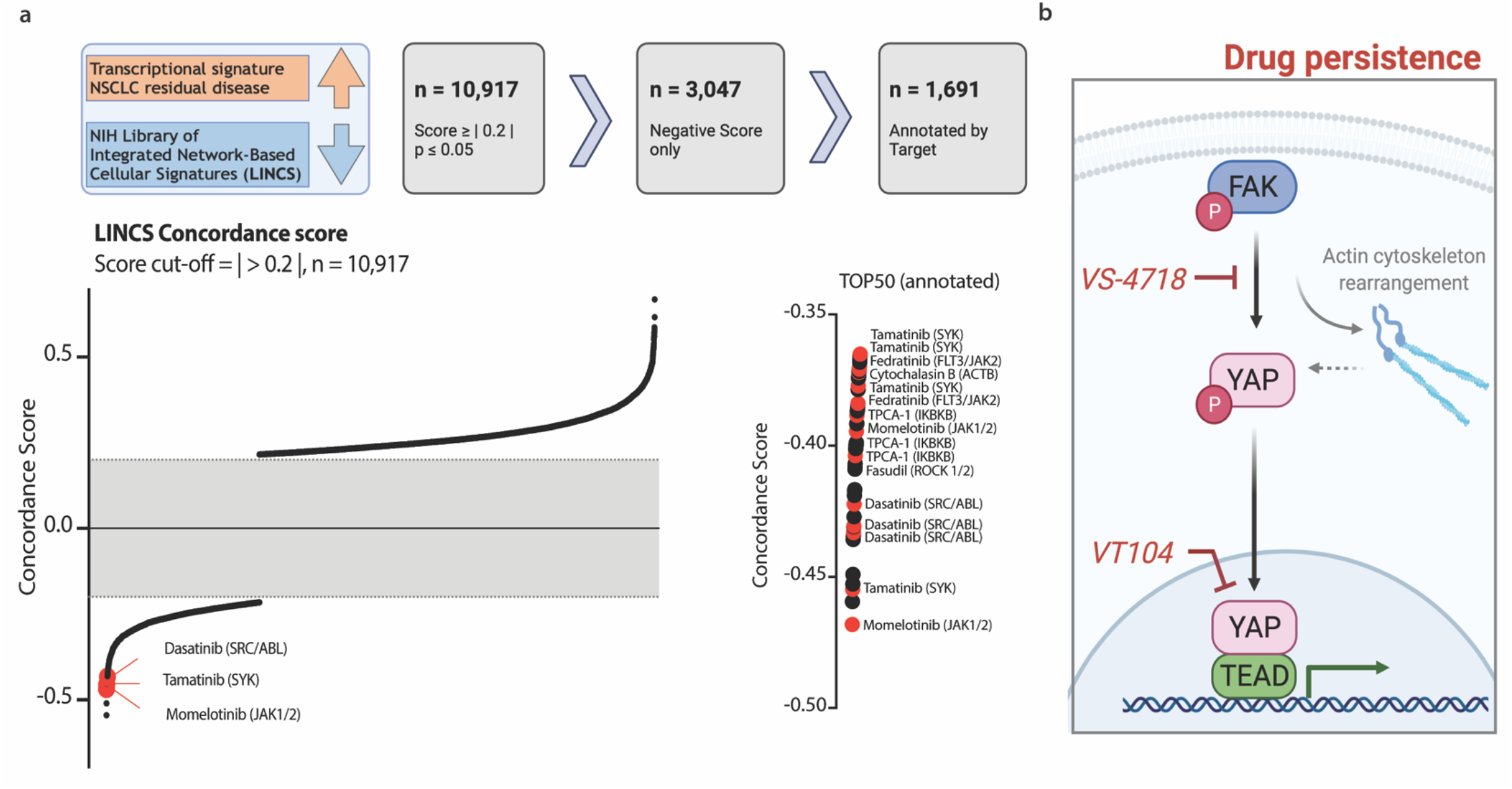
Correlation of drug-mediated transcriptional changes and expression profiles at residual disease. (**a**) Concordance score for the correlation of transcriptional scRNAseq profile at the RD timepoint and LINCS L1000 data collected for drug-mediated expression changes. Schematic of data processing (top) as well as distribution of concordance scores (bottom) are presented. Sigmoidal distribution of concordance scores across drug perturbations showing significant correlation with residual disease-associated expression changes (*n* = 10,917). Top 50 negative concordance scores are presented in detail, with inhibitors targeting relevant proteins for focal adhesion / SRC or inflammatory signaling highlighted in red and annotated. Schematic diagram was created with BioRender.com. (**b**) Pathway schematic for relevant changes in drug-tolerant persister cells, highlighting mechanistic nodes of YAP-mediated transcriptional adaptation and therapeutic vulnerabilities. Schematic diagram was created with BioRender.com.

## DISCUSSION

The preclinical and clinical data we present highlight the relevance of FAK signaling for YAP nuclear localization and drug persistence upon treatment with targeted inhibitors (Fig. 8b). We reveal YAP-driven transcriptional adaptation as a functional mechanism in drug tolerance, with FAK signaling critically involved in mediating YAP nuclear relocalization. Experiments in a humanized mouse model confirmed the role of YAP in reducing treatment sensitivity to targeted inhibitors and indicate a tumor microenvironment supporting tumor outgrowth. Further, therapeutic vulnerabilities targeting residual cells are present mechanistically upstream and downstream of YAP and include FAK inhibition by VS-4718 as well as YAP/TEAD inhibition by VT104 (Fig. 8b). Clinical relevance of the highlighted mechanisms and YAP activity in residual disease were shown by the analysis of unique clinical specimens and a chemical-genetics large scale dataset analysis to identify drug-based perturbations that highlight potential dependencies that can be exploited to target residual disease, including but not limited to the FAK-YAP/TEAD signaling axis.

Clinical approaches targeting the FAK-YAP/TEAD signaling axis are emerging, with recent efforts focused on the development of TEAD inhibitors^13, 40, 47^. Inhibition of TEAD S-palmitoylation by small molecules has been reported to impact TEAD function and to block YAP-TEAD interaction^47^. Optimization of TEAD palmitoylation inhibitors has overcome pharmacological limitations and resulted in *in vivo* efficacy for VT104 in *NF2*-deficient mesothelioma xenografts^40^. This work provided a rational for the initiation of a phase I clinical trial (NCT04665206)^40, 48^. Similarly, targeting FAK signaling has gained recent momentum, in particular as a combinatorial agent overcoming resistance-associated signaling in cancer therapy^49^. Amongst other contexts, the engagement of FAK signaling has been reported in KRAS-mutant patients treated with the dual RAF/MEK inhibitor VS-6766 in a phase I clinical trial (NCT03875820) ^49–51^, with subsequent and current clinical studies testing the efficacy of combinatorial treatment with FAK and RAF/MEK inhibitors (NCT04620330, NCT04625270) ^52, 53^. Our findings provide evidence of FAK-YAP/TEAD signaling engagement in drug tolerance across EGFR-mutant, ALK fusion-positive, KRAS-mutant and NF1-mutant NSCLC more broadly and therefore offer rationale for combinatorial treatment approaches testing FAK inhibitors (e.g., VS-4718) and TEAD inhibitors (e.g., VT104) across several molecularly defined NSCLC subtypes to enhance treatment response to targeted inhibitors against these important oncogenic targets.

FAK inhibitors have shown promising anti-tumor immune effects and combinatorial treatment of FAK and immune checkpoint inhibition is currently being evaluated in a phase I/IIa clinical study across pancreatic cancer, mesothelioma, and NSCLC (NCT02758587)^49, 54^. Our prior work showed significant immune cell alterations at residual disease, characterized by higher T cell infiltrates and continued presence of pre-dysfunctional cytotoxic CTLA4-positive T cells^7, 55^. The FAK-associated tumor-supportive immune microenvironment changes indicate a possible benefit of combinatorial treatment with FAK inhibitors regarding the normalization of an effective immune response at residual disease. Similarly, YAP/TEAD-mediated microenvironmental changes need to be further studied in the respective immune competent mouse models and could result in additional clinical avenues.

Overall, our findings provide evidence for a distinct mechanism of drug tolerance and residual disease centered on FAK-YAP/TEAD signaling axis activation that promotes cancer cell survival and potentially tumor immune evasion during targeted therapy. This study highlights a targetable pathway (FAK-YAP/TEAD) that limits response to current targeted therapies in a relatively conserved manner in oncogene-driven NSCLCs and the potential importance of biological crosstalk between cancer cell signaling events (e.g., FAK-YAP) and the TME. Our findings have implications for developing improved treatment regimens to thwart tumor progression on therapy by targeting residual disease to enhance clinical responses in oncogene-driven NSCLC patients.

## ONLINE METHODS

### Cell lines and culture reagents

PC9 and derived isogenic cell lines, NCI-H1975 and derived isogenic cell lines, NCI-H3122, NCI-H2228, STE- 1, NCI-H358, and NCI-H1838 cells were grown in RPMI medium 1640 supplemented with 10% (v/v %) FBS, 100 IU/mL penicillin, and 100 μg/mL streptomycin. 293T cells were grown in Dulbecco’s Modified Eagle Medium (DMEM) supplemented with 10% (v/v %) FBS, 100 IU/mL penicillin, and 100 μg/mL streptomycin. All cells were maintained at 37 °C in a humidified atmosphere at 5% CO2.

### Persister cell generation

For persister cell generation and subsequent analysis by immunoblotting or RNA sequencing, 5x10^5^ cells were seeded in 10 cm culture dishes. For persister cell generation and subsequent analysis by confocal microscopy or PLA assay, 5x10^4^ cells were seeded in 35mm glass-bottom dishes (MatTek Corporation). After incubation overnight, cells were treated with an IC80 concentration of the respective targeted inhibitor. Treatment was replenished every 3-4 days. Persisters cells were harvested as indicated and after ≥ 7 days of drug exposure. For comparison, parental cells were treated for 48 h with 0.1% DMSO (DMSO) or with an IC80 concentration of the respective targeted inhibitor (ACUTE). Acquired resistant cell lines were maintained in the drug directly after seeding and treatment was replenished for 48 h with an IC80 concentration of the respective targeted inhibitor (AR or RESISTANT).

### Antibodies

For Western blotting, antibodies for phospho-ACK1 (Y284, #3138), Bcl-xL (#2764), phospho-EphB1 (Y324, #3481), EphB1 (#3980), ErbB2 (#4290), ErbB3 (#4754), phospho-FAK (Y397, #8556), FAK (#3285), FGFR1 (#9740), Histone H3 (#9715), Lamin B1 (#12586), phospho-LATS1 (T1069, #8654), LATS1 (#3477), phospho-SRC (Y416, #2101), SRC (#2108), phospho-YAP (S127, #13008), and YAP/TAZ (#8418) were purchased from Cell Signaling Technology. The antibody for phospho-YAP (Y357, #62751) was purchased from Abcam. Antibodies for ACK1 (#sc-28336), FGFR2 (#sc-6930), and GAPDH (#sc-365062) were purchased from Santa Cruz Biotechnology. The antibody for β-actin (#A2228) was purchased from Sigma-Aldrich. Antibodies were diluted according to the manufacturer’s recommendations. For confocal analysis, PLA, and immunoprecipitation, pan-TEAD (#13295) and YAP (#12395) antibodies were purchased from Cell Signaling Technologies and diluted 1:100. For immunohistochemistry, the antibody for YAP (clone H-125, #sc-15407) was purchased from Santa Cruz Biotechnology and diluted 1:150. For immune cell profiling in humanized mice models, fluorochrome-conjugated monoclonal antibodies to the following human antigens were used: CD45-Alexa Fluor 700 (clone 2D1, HI30), CD45-phycoerythrin (PE; clone 2D1, HI30), CD3-PerCp/cy5.5 (clone HIT3a), CD19-PE-cyanine 7 (clone HIB19), CD8-allophycocyanin-cyanine 7 (clone RPA-T8, HIT8a), CD4-Pacific blue (clone OKT4), HLA-DR-PerCp/cy5.5 (clone LN3), CD11b-PE-Cy7 (clone 1CRF-44) (Thermo fisher), CD25-APC (clone CD25-4E3), CD163-APC (clone ebioGH1/61; Thermo fisher). A mouse CD45-FITC (clone 30-F11) antibody was used for gating out murine leukocytes. Most antibodies were purchased from Biolegend, if not otherwise mentioned.

### Pharmacologic agents

Osimertinib (AZD9291), alectinib (CH5424802), ARS-1620, trametinib (GSK1120212), VS-4718 (PND-1186), and dasatinib were purchased from Selleck Chemicals. The TEAD inhibitor VT104 was kindly provided by Vivace Therapeutics, Inc. The SHP2 inhibitor RMC-4550 was kindly provided by Revolution Medicines, Inc..

### High-content microscopy screening

Cell lines were seeded in 96-well assay clear-bottom microplates at a density of 2,500-5,000 cells per well in a total volume of 90 μL per well and incubated at 37 °C, 5% CO2 overnight. Following drug exposure, cell confluency was measured by staining with Hoechst 33342 (Thermo Fisher Scientific) nuclear dye; apoptosis was measured using YO-PRO-1 early apoptosis dye (Thermo Fisher Scientific) and analyzed using a CellInsight High-Content Microscope (Thermo Fisher Scientific) at the indicated time points.

### Apoptosis analysis

Apoptotic cell death was detected by flow cytometry using Annexin V and 7-amino-actinomycin (7-AAD) staining. Cells were harvested and resuspended in Annexin V-binding buffer containing 10% Annexin V-FITC and 10% 7-AAD staining solution (Thermo Fisher Scientific). After an incubation time of 15 min at 4°C, stained cells were analyzed by flow cytometry.

### Western blot analysis

Whole-cell lysates were prepared by using radio-immunoprecipitation assay buffer (RIPA) [10 mM Tris·Cl (pH 8.0), 1 mM EDTA, 0.1% sodium deoxycholate, 0.1% SDS, 140 mM NaCl] supplemented with protease inhibitor and phosphatase inhibitor (Roche). Nuclear-cytoplasmic extracts were prepared using 0.1% NP-40 in PBS supplemented with protease inhibitor and phosphatase inhibitor (Roche) as previously described^56^. Whole-cell and nuclear lysates were clarified by centrifugation at 17,000 x g for 15 minutes at 4 °C. Lysates were quantified using the Pierce BCA Protein Assay Kit (Thermo Fisher Scientific). Equal masses of protein (5 - 20 ug) were separated by 4–15% of SDS/ PAGE and were transferred onto nitrocellulose membranes (Bio-Rad) for protein blot analysis. After blocking in 5 % milk/ Tris-buffered saline, 0.1% Tween-20 (TBS-T), membranes were incubated with primary antibody overnight at 4 °C, then washed and incubated with secondary antibody for 1 hour at room temperature. Protein bands were visualized using either a fluorescence system (LI-COR) or Amersham ECL chemiluminescent reagent (GE Life Sciences); chemiluminescent signals were visualized with an ImageQuant LAS 4000 instrument (GE Healthcare).

### Generation of endogenously tagged YAP-mNeonGreen2 cell lines

Generation of endogenously tagged mNeonGreen21-10/11 cell lines was performed in EGFR-mutant PC9 cells as described previously^57^ using the sgRNA spacer sequence 5’-AGGCAGAAGCCATGGATCCC-3’. Isogenic cell lines (1-E7 and 2-G10) were generated by single-cell sorting via fluorescence-activated cell sorting (FACS) and outgrowth to stable cell lines. Integration of mNeonGreen211 was confirmed by genomic sequencing and by a reduction in fluorescence upon gene knockdown. Isogenic cell lines showed osimertinib responses equal to parental bulk PC9 cells as evaluated by CellTiter-Glo assay.

### Confocal analysis

Cells were seeded in 35mm glass-bottom dishes (MatTek Corporation) or µClear 96-well imaging plates (Greiner Bio-One). At the time of harvest, cells were washed carefully and fixed in 4% paraformaldehyde. Cells were then permeabilized in 0.1% Triton-X/PBS and blocked in 5% bovine serum albumin (BSA) / 0.1% Triton-X/PBS. Primary antibody was diluted in 1% BSA / 0.1% Triton-X/PBS and incubated overnight at 4 °C. After washing, secondary antibodies were diluted in 1% BSA / 0.1% Triton-X/PBS and added for 1 hour at room temperature. Where indicated, actin filaments were stained with rhodamine-phalloidin (Thermo Fisher Scientific) as described by the manufacturer. After secondary antibody staining, cells were washed and then stained with DAPI solution (1:1000 in PBS, stock 1 mg/mL, Thermo Fisher Scientific). For endogenously tagged YAP-mNeonGreen2 cells, permeabilization, blocking as well as primary and secondary antibody staining were omitted. Cells were imaged at a Yokogawa CSU22 spinning disk confocal microscope using a Plan Apo VC 60X/ 1.4 Oil objective (Nikon Imaging Center, UCSF). Image analysis was done via Fiji ImageJ software^58^. Quantification of relative integrated density for nuclear levels was performed by automated analysis quantifying the intensity for the protein of interest per nuclei.

### PLA assay

Cells were seeded in 35mm glass-bottom dishes (MatTek Corporation). Proximity ligation assays were performed using the Duolink In Situ Red Starter Kit Mouse/Rabbit (Millipore Sigma). In brief, cells were washed carefully and fixed in 4% paraformaldehyde. Cells were then permeabilized in 0.1% Triton-X/PBS before blocking. Primary antibodies were added overnight at 4 °C. PLA probes were added for 1 hour at 37 °C before ligation and amplification. After washing and staining cell nuclei with DAPI solution (1:1000 in PBS, stock 1 mg/mL, Thermo Fisher Scientific), cells were imaged at a Yokogawa CSU22 spinning disk confocal microscope using a Plan Apo VC 60X/ 1.4 Oil objective (Nikon Imaging Center, UCSF). Image analysis was done via Fiji ImageJ software^58^ and PLA signals per nuclei were counted.

### Endogenous immunoprecipitation

Primary antibodies were coupled to Dynabeads Protein G beads at a 1:5 ratio (antibody:beads, v/v) by constant rotation for 6 hours at 4 °C. Nuclear fractions of persister cells were prepared using the NE-PER Nuclear and Cytoplasmic Extraction Reagents (Thermo Fisher Scientific) and kept on ice. Lysates were quantified using the Pierce BCA Protein Assay Kit (Thermo Fisher Scientific). Equal masses of proteins (300 µg) were added to antibody-coated beads and incubated by constant rotation overnight at 4 °C. Bead-coupled samples were washed and resuspended in 4x Laemmli buffer [277.8 mM Tris-HCl, pH 6.8, 44.4% (v/v) glycerol, 4.4% LDS, 0.02% bromophenol blue, supplemented with 10% (v/v) 2-Mercaptoethanol]. Beads were collected and samples were analyzed by Western blot analysis as outlined above.

### Knock-down and CRISPR knock-out

CRISPR-mediated YAP and FAK knock-out cells were engineered by Synthego (Synthego Corporation, Redwood City, USA). Transient YAP silencing was achieved by knock-down using individual Dharmacon ON-TARGETplus YAP1 siRNAs (siYAP#1: J-012200-08, siYAP#2: J-012200-07) compared to non-target control (D-001810-02). Knock-down of focal adhesion kinase signature genes EphB1 (siEPHB1: L-003121-00), FAK (siPTK2B: L-003165-00), and ACK1 (siTNK2: L-003102-01) as well as of LATS1 (L-004632-00) and LATS2 (L-003865-00) was induced using SMARTpool Dharmacon ON-TARGETplus siRNAs compared to non-target control (D-001810-10). Target cells were transiently transfected using Lipofectamine RNAiMAX Transfection Reagent (Thermo Fisher Scientific). SiRNA-mediated knock-down was initiated either at the beginning of treatment with the targeted inhibitor (knock-down during persister cell generation) or when persisters cells were established after 7 days of treatment (knock-down at persister cell state). In both cases, knock-downs were repeated every three days and a consecutive number of three knock-downs was performed before harvest. Knock- out and knock-down of the protein of interest were verified by Western blot analysis.

### YAP-5SA overexpression

Full-length YAP was amplified by PCR using forward primer 5’- TTTGACCTCCATAGAAGATTCTAGATGGAACAAAAACTCATCTC-3’ and reverse primer 5’- AGCGATCGCAGATCCTTCGCGGCCGCTATAACCATGTAAGAAAGCTTTC-3’ from pQCXIH expression constructs encoding myc-tagged YAP-WT (Addgene #33091), YAP-5SA (Addgene #33093), and YAP-S94A (Addgene #33094), respectively. After XbaI/NotI digestion, PCR products were cloned into the lentiviral pCDH-puro plasmid backbone and correct insertion was verified by Sanger sequencing. For lentivirus production, 293T cells were co-transfected with pCDH-YAP expression plasmids and lentiviral packaging plasmids pCMV-dR8.91 and pMD2.G using the TransIT-LT1 Transfection Reagent (Mirus Bio). Viral supernatant was harvested 72 hours after transfection. Target cells were infected and selected with 1 µg/mL puromycin. YAP overexpression and YAP nuclear localization upon treatment with 0.1% DMSO or 2 µM osimertinib in stable transduced PC9 cells was monitored by Western blot analysis and confocal microscopy.

### Cell viability and persister cell numbers

Cell survival and persister cell numbers upon genetic or pharmacologic perturbations were evaluated by counting cells using a Vi-CELL XR Cell Viability Analyzer (Beckman Coulter, Inc.). Response to escalating drug doses in stable transduced PC9 and H358 cells was analyzed by CellTiter-Glo assay (Promega). For the latter, cells (5x10^3^/ well) were seeded in clear-bottom 96-well plates. After overnight incubation, cells were treated with escalating drug concentrations and harvested at day 5 post drug treatment.

### RNA sequencing and gene set enrichment analysis

RNA was extracted from snap-frozen tissue or cell pellets. For tissue samples, tissue was minced using a liquid nitrogen-cooled mortar and pestle before RNA extraction. RNA isolation was performed using the RNeasy Mini kit (Qiagen) including an on-column DNase I digestion. RNA quality was assessed by automated electrophoresis using the RNA 6000 Pico Kit and an Agilent 2100 BioAnalyzer (Agilent Technologues, Inc.). RNA was quantified using the Qubit RNA HS Assay Kit and a Qubit 2.0 fluorometer (Thermo Fisher Scientific). Library preparation and paired-end 150bp (PE150, Illumina) RNA sequencing was performed by Novogene (Novogene Corporation, Sacramento, USA). RNA-Seq reads were mapped to the hg19 reference genome using STAR (Spliced Transcripts Align to a Reference, v2.4.2a). The expression level of transcript per million (TPM) reads were quantified using RNA-Seq by Expectation-Maximization algorithm (RSEM v1.2.29). The quantified gene expressions of 26,334 transcripts (including coding genes and non-coding genes) were processed in R studio. Differentially expressed genes between tumor and normal samples were identified using the EdgeR algorithm. Gene set enrichment analysis was done using GSEA 4.0.1 software^59, 60^.

### scRNA sequencing trajectory

A BFP-tagged barcode library (Addgene #85968) was lentivirally infected into isogenic EGFR-mutant PC9-C2 and H1975-B10 cells. Cells were sorted and serially titrated to allow for ∼1000 unique lineages. After expansion, cells were subjected to 0.1% DMSO or 2 µM osimertinib treatment and frozen down at the indicated timepoints. Cells were thawed, hashed with TotalSeq A anti-human hashtag antibodies (Biolegend), and pooled for single- cell RNA sequencing on the 10X chromium v3 platform (10x Genomics). Cell hash libraries were prepared as specified by Biolegend. Custom barcode amplification was performed by two rounds of PCR. Libraries were sequenced on the NovaSeq Illumina platform (Center for Advanced Technology, UCSF). After NGS sequencing, cells were called with 10X Cell Ranger pipeline and cell hashes were called using the scEasyMode package in Python. In addition, bulk genomic barcodes were prepared from the same time points used for single-cell RNA sequencing using the Quick Extract gDNA extraction protocol (Lucigen Corporation) and custom barcode amplification primers for NGS library preparation. A custom script for calling genomic barcodes mapping between single-cell genomic barcodes and bulk genomic barcodes collected from the same samples was used to assess population frequency and map onto single-cell transcriptomes. The diversity index was calculated as 1 - Sum_i (pi^2), where pi is the relative abundance of line i. The diversity index is at its maximum when all lineages are equally abundant and decreases if some lineages are enriched and others depleted. The index was scaled by the max possible index given the number of lineages which is max(Lineage diversity index) = 1 - n[(1/n)^2] = 1 - 1/n; n: number of lineages.

### Whole exome sequencing

DNA was extracted from snap-frozen cell pellets using the DNeasy Blood & Tissue kit (Qiagen). DNA quality was assessed by automated electrophoresis using the High Sensitivity DNA Kit and an Agilent 2100 BioAnalyzer (Agilent Technologies, Inc.). DNA was quantified using the Qubit dsDNA HS Assay kit and a Qubit 2.0 fluorometer (Thermo Fisher Scientific). Library preparation and paired-end 150bp (PE150, Illumina) DNA sequencing were performed by Novogene (Novogene Corporation, Sacramento, USA). Pair-end fastq files were mapped to the hg19 genome and mutation calling using the SeqMule pipeline^61^. The vcf files were annotated using ANNOVAR software at a high-performance computing cluster (UCSF Helen Diller Comprehensive Cancer Center). Further analysis of annotated variants was conducted under the RStudio/R environment.

### Patient samples and use of human tissue

All patients gave informed consent for collection of clinical correlates, tissue collection, research testing under Institutional Review Board (IRB)-approved protocols. Patient studies were conducted according to the Declaration of Helsinki, the Belmont Report, and the U.S. Common Rule.

### NSCLC organoid cultures

Organoid cultures from NSCLC specimens were established as previously described^62, 63^. Organoid cultures were embedded in Reduced Growth Factor Basement Membrane Extract, Type 2 (BME2) matrix (Thermo Fisher Scientific) and maintained in Dulbecco’s Modified Eagle’s Medium/Ham’s nutrient mixture F12 (DMEM/F-12) GlutaMAX supplement, supplemented with 100 U/mL penicillin/streptomycin, 10 mM HEPES, 25 nM hRspondin, 1x B27, 5 mM Nicotinamide, 1.25 mM N-Acetylcysteine, 500 nM A-8301, 500 nM SB202190, 50 µg/mL Primocin, 100 ng/mL hNoggin, 100 ng/mL hFGF-10, and 25 ng/mL hFGF-7^63, 64^. Mutational profiling of organoid cultures was performed by whole-exome sequencing. Signaling alterations upon osimertinib treatment in EGFR-mutant organoids were evaluated by treating single suspensions for 2 hours and subsequent Western blot analysis. Drug sensitivity was analyzed by 3D CellTiter-Glo assay (Promega). In brief, single cells suspensions were prepared by TrypLE digestions, and cells (7.5x10^3^/ well) were seeded in BME2 on clear-bottom 96-well plates (Corning). After seven days in culture, newly formed organoids were treated with indicated drug concentrations in reduced growth factor media. Five days after treatment initiation, the viability of cells was assessed. Persister cell generation of organoids was performed as outlined, seeding NSCLC organoid cells (1.8x10^5^/ well) embedded in BME2 in a 6-well plate format. After three days in culture, organoids were treated with 0.1% DMSO or 1 µM osimertinib. DMSO-treated control cells were harvested three days after treatment. For persisters, the drug was replenished every 3 days, and cells were harvested post 11 days on treatment. The engagement of YAP signature genes was evaluated by RNA sequencing and gene set enrichment analysis.

### Subcutaneous xenograft and PDX experiments

All animal experiments were conducted under UCSF IACUC-approved animal protocol no. AN187306-01B. H1975 tumor xenografts were generated by injection of one million cells in a 50/50 suspension of matrigel/PBS into 6- to 8-wk-old female SCID mice. Once the tumors grew to an average size of ∼200 mm^3^, mice were randomized and treated with vehicle (2% HPMC E-50, 0.5% Tween-80 in 50 mM Sodium Citrate Buffer, pH 4.0), 5 mg/kg osimertinib q.d., 50 mg/kg VS-4718 b.i.d., 3 mg/kg VT104 q.d. as well as combinations of osimertinib with VS-4718 or VT104. No substantial toxicity was observed in mice treated with either combination regimen incorporating FAK or TEAD inhibitors by assessment of body weight (Supplementary Fig. 8c) and general animal well-being. EGFR-mutant TH021 and ALK fusion-positive LG0812 PDX, tumors were propagated into 6- to 8-wk-old female SCID mice. Once the tumors grew to an average size of ∼400 mm^3^, mice were randomized and treated with vehicle, 10 mg/kg osimertinib q.d. (TH021) or 6 mg/kg alectinib q.d. (LG0812). For the EGFR-mutant TH021 PDX model, combinatorial treatment effects were assessed upon treatment with vehicle, 5 mg/kg osimertinib q.d., 3 mg/kg VT104 q.d. as well as combinations of osimertinib with VT104. Tumor volume was assessed regularly. At the treatment endpoint, tumors were halved and harvested in ice-cold PBS. One-half of tumors were incubated in 10% neutral-buffered formalin for 24-72 hours and then stored in 70% Ethanol until embedding in paraffin blocks and subsequent analysis by immunohistochemistry. The other half was snap-frozen for later analysis by RNA sequencing.

### Humanized mouse model

Humanized xenograft models were kindly established by the laboratory of Jack Roth at MD Anderson Cancer Center as described previously^42^. All animal use was conducted in accordance with the guidelines of the Animal Care and Use Committee of MD Anderson Cancer Center. In brief, female 3-to-4-week-old NOD.Cg-Prkdcscid Il2rgtm1Wjl/SzJ (NSG) mice, which are suitable for the engraftment of human hematopoietic cells), were housed in microisolator cages under specific pathogen-free conditions in a dedicated humanized mice room in the animal facility at The University of Texas MD Anderson Cancer Center. Mice were given autoclaved acidified water and fed a special diet (Uniprim diet). Human umbilical cord blood units were obtained from MD Anderson Cord Blood Bank under an IRB-approved protocol. Fresh cord blood units were delivered within 24 h of harvest and were HLA typed immediately at MD Anderson HLA-typing core facility. Cord blood was diluted to a ratio of 1:3 with phosphate-buffered saline, and mononuclear cells were isolated by using density-gradient centrifugation on Ficoll medium. CD34+ HSPCs were isolated using a direct CD34+ MicroBead kit (Miltenyi Biotec). NSG mice were irradiated with 200 cGy using a 137Cs gamma irradiator. Over 90% pure freshly isolated CD34+ HSPCs were injected intravenously, 24 h after irradiation, at a density of 1 to 2 × 105 CD34+ cells/ mouse. All Hu-NSG mice were verified for humanization before tumor implantation. For PC9-parental, PC9-WT (YAP overexpression), PC9-5SA (hyperactive YAP) cell lines, 5-7x106 cells were injected subcutaneously 8-weeks post humanization of mice. Another 3-4 weeks post tumor cells implantation in humanized mice and when tumor sizes reached 200mm3, animals were randomized into treatment and no-treatment groups based on tumor size and donor HLA type. Five mice per group from multiple umbilical cord blood donors were used. Mice were treated with vehicle or osimertinib (5 mg/kg) orally 5 days a week for consecutive 3 weeks. For immune analysis, erythrocytes in the peripheral blood were lysed with ACK lysis buffer (Fisher Scientific). Single-cell suspensions were prepared. Several 10-color flow cytometry panels were used for immune profiling of both innate and adaptive immune populations in humanized mice and for evaluating immune response after treatment. All samples were run on Attune NxT flow cytometer (Thermo fisher), and data were analyzed by Flow Jo and Kaluza software packages.

### Immunohistochemistry

Formalin-fixed paraffin-embedded (FFPE) tumor blocks were cut at 4-micron thickness and mounted as sections on positively charged histology slides. Immunohistochemistry staining was performed as described previously^65^. In brief, slides were deparaffinized in xylene, rehydrated and epitope retrieval was induced in a histology pressure cooker using pH 6.1 citrate buffer (Dako Denmark A/S, S2369). After endogenous peroxidase and protein block, slides were incubated with primary antibody solution overnight at 4 °C. Then slides were incubated with secondary antibody for 30 minutes (EnVision Dual Link Labelled Polymer HRP, Agilent K4065), stained using 3,3-DAB, and counterstained with hematoxylin. Slides were dehydrated and mounted before digitization using an Aperio AT2 Slide Scanner (Leica Biosystems) at a 20X objective. Quantification of nuclear YAP levels was performed via the Aperio Image Scope digital pathology software using the nuclear quantification algorithm.

### YAP signature in scRNA seq data of patient samples

Single-cell sequencing data were derived from previously published work^7^ and filtered to limit analysis to malignant lung epithelial cells only (*N* cells: TN = 621, RD = 484, PD = 138). Differentially expressed gene sets for YAP activation (YAP-5SA-UP), persistence (PC9, H3122, H358 and overlapping genes across all cell line models [overlap persisters]), and patient treatment timepoint (TN, RD, PD) were compared to the patient gene expression dataset via permutation analysis (R package, GSALightning, v.1.1.7). Subject classes were assigned to every single cell based on the corresponding time point, where RD = “RD” and TN or PD = “nonRD”. Each gene was tested for significance with unpaired t-tests and the mean across genes was used to calculate the gene set statistics. Multiple testing correction was done via Benjamini-Hochberg. The union of significantly upregulated genes from the YAP-5SA_UP gene set, significantly upregulated genes across persister models, and the RD treatment timepoint were then used to define the YAP gene signature (adjusted *p*-value < 0.05). Other signatures (cell cycle, FAK) were determined from relevant literature^7, 31^. Signature expression for single-cell data was processed and plotted in R with ggplot2 (v.3.3.3) and Seurat (v.3.2.2).

### Survival analysis of YAP gene signature

TCGA LUAD data was retrieved (https://portal.gdc.cancer.gov/projects/TCGA-LUAD). Samples were grouped according to their expression quantile (High > 0.75; 0.25 ≤ Medium ≥ 0.75; Low < 0.25) for the sum gene expression of the corresponding signature. Correlation of signature expression to patient overall survival was assessed. For the latter, data were processed in R with survival (v.3.2) using default arguments for the single-outcome, single-event type and depicted via Kaplan-Meier (log-rank test) curves.

### LINCS L1000 concordance score

The NIH LINCS L1000 database contains gene expression data from cultured human cells that were treated with small molecule and genetic perturbagens. Level 4 data was sourced from the Gene Expression Omnibus Series GSE70138. Expression data was restricted to small-molecule perturbagens and overlapped with the residual disease signature (*N* = 83 genes). Using a previously published computational pipeline^66, 67^ (*65*, *66*), a score for each signature-drug pair was determined using a non-parametric rank-based method that is similar to the Kolmogorov–Smirnov test statistic, where negative scores indicate genes in the ranked drug profile are oppositely regulated in the ranked signature. For the upregulated NSCLC residual disease signature, drug-gene expression profiles were chosen that produce the greatest significant negative score. *P*-values for drug profiles were determined by comparing their scores to a distribution of random scores and adjusted with the false discovery rate (FDR; Benjamini-Hochberg, *α* = 0.05) method.

### Statistical analysis

Quantitative data are presented as mean +/− standard deviation (S.D.). Statistical tests were performed using GraphPad Prism 8.4.2. Two-sided Student’s *t*-tests were used for comparisons of the means of data between two groups unless otherwise specified. For comparisons among multiple independent groups, a one-way ANOVA test was used. For animal studies, animals were randomized before treatments, and all animals treated were included for the analyses.

## SUPPLEMENTARY INFORMATION

Supplementary Fig. 1. Characterization of drug-tolerant persister cells.

Supplementary Fig. 2. Signaling and transcriptional changes in drug-tolerant persister cells.

Supplementary Fig. 3. YAP as a functional marker in drug-tolerant persister cells.

Supplementary Fig. 4. Changes in drug tolerance upon YAP knock-down or overexpression.

Supplementary Fig. 5. Single cell RNA sequencing trajectory.

Supplementary Fig. 6. Sensitivity to combinatorial treatment modalities.

Supplementary Fig. 7. Treatment response in PDO and PDX models.

Supplementary Fig. 8. Combinatorial treatment studies in EGFR-mutant H1975 xenograft model.

Supplementary Fig. 9. Combinatorial treatment with TEAD inhibitor VT104.

Supplementary Fig. 10. Combinatorial treatment studies in EGFR-mutant TH021 PDX model.

Supplementary Fig. 11. Transcriptional changes and YAP nuclear localization in patient specimens.

Supplementary Fig. 12. LINCS L1000 concordance analysis.

## Acknowledgements

The authors would like to acknowledge Dana S. Neel, Manasi K. Mayekar, Beatrice Gini, Nilanjana Chatterjee, Ross A. Okimoto, Anatoly Urisman, and Johannes R. Kratz for their scientific input and experimental help. This research project was conducted with support from National Institutes of Health (NIH)/National Cancer Institute (NCI): U54CA224081 (T.G.B., C.J.K.) and U54 DRSN supplement (J.A.R.),The University of Texas MD Anderson Cancer Center’s Cancer Center Support Grant (CCSG) CA-016672 - Lung Program and Shared Core Facilities (J.A.R.), Specialized Program of Research Excellence (SPORE) Grant CA- 070907 (J.A.R.), PDX development and trial grant U54CA-224065 (J.A.R.), Lung Cancer Moon Shot Program (J.A.R.), U01 grants: U01CA217882 (T.G.B.), U01CA217851 (C.J.K.), R01 grants: R01CA231300 (T.G.B., B.H.), R01CA204302 (T.G.B.), R01CA211052 (T.G.B.), R01CA169338 (T.G.B.), R01-CA18731 (J.P.R.), R01GM131641 (B.H.); NIH/National Institute of Allergy and Infectious Diseases (NIAID): R01-AI104789 (J.P.R.); NIH/National Heart, Lung, and Blood Institute (NHLBI): R01 - HL120724 (J.P.R.); The University of Texas MD Anderson Cancer Center, sponsored research agreement from Genprex, Inc. (J.A.R.), UCSF PBBR TMC (Technologies, Methodologies, and Cores) grant, gift from the UCSF Pathology department (J.P.R.); Mark Foundation for Cancer Research, Endeavor Program grant A136299 (J.P.R.); The Ludwig Cancer Foundation (C.J.K.); Stand Up To Cancer Foundation (C.J.K.); The Damon Runyon Cancer Research foundation, P0528804 (C.M.B.); Doris Duke Charitable Foundation P2018110 (C.M.B.); V Foundation P0530519 (C.M.B.); The Van Auken Foundation and Addario Lung Cancer Foundation, Young Innovators Team Award (J.W.R.); Ministerio de Ciencia e Innovacion, Spain, PID2019-105303RB-I00/AEI/10.13039/501100011033 (P.S.); Comunidad de Madrid, Spain, B2017/BMD-3724 (P.S.); Fundación Española Contra el Cáncer (AECC), Spain, GCB14142311CRES (P.S.); FPI fellowship from Universidad Autónoma de Madrid, Spain (C.F.M.); Travelling Fellowship from The Company of Biologists (C.F.M.); the German Cancer Aid, Mildred Scheel postdoctoral fellowship (F.H.). B.H. is a Chan Zuckerberg Biohub Investigator.

## Author contribution

F.H. and T.G.B. designed the study. F.H., C.F.M., L.C., I.M.M., J.Y., V.O., D.B.R., C.G., D.L.K., D.V.A., J.G., K.N.S., K.A.H., O.M.G., W.T., J.K.R., and W.W. performed experiments, collected, and analyzed data. C.E.M., J.W.R., and C.M.B. coordinated the availability of clinical specimens. M.M., J.S.G., C.J.K., J.P.R., T.T.T., L.P., B.H., H.G., P.S., S.B., W.W., and C.M.B. provided scientific input. J.A.R. provided scientific input and oversaw experiments. F.H. and T.G.B. wrote the manuscript. T.G.B. oversaw the study. All authors have approved the manuscript.

## Data and materials availability

This data is available as an NCBI Bioproject under accession number PRJNA766057. For single cell RNA seq analyses of patient specimens, the data is derived from a previously published study ^68^ and available as an NCBI Bioproject under accession number PRJNA591860. Plasmids, data and codes generated are available by request from the corresponding author.

## SUPPLEMENTARY INFORMATION

### Supplementary Figures

**Supplementary Fig. 1.**
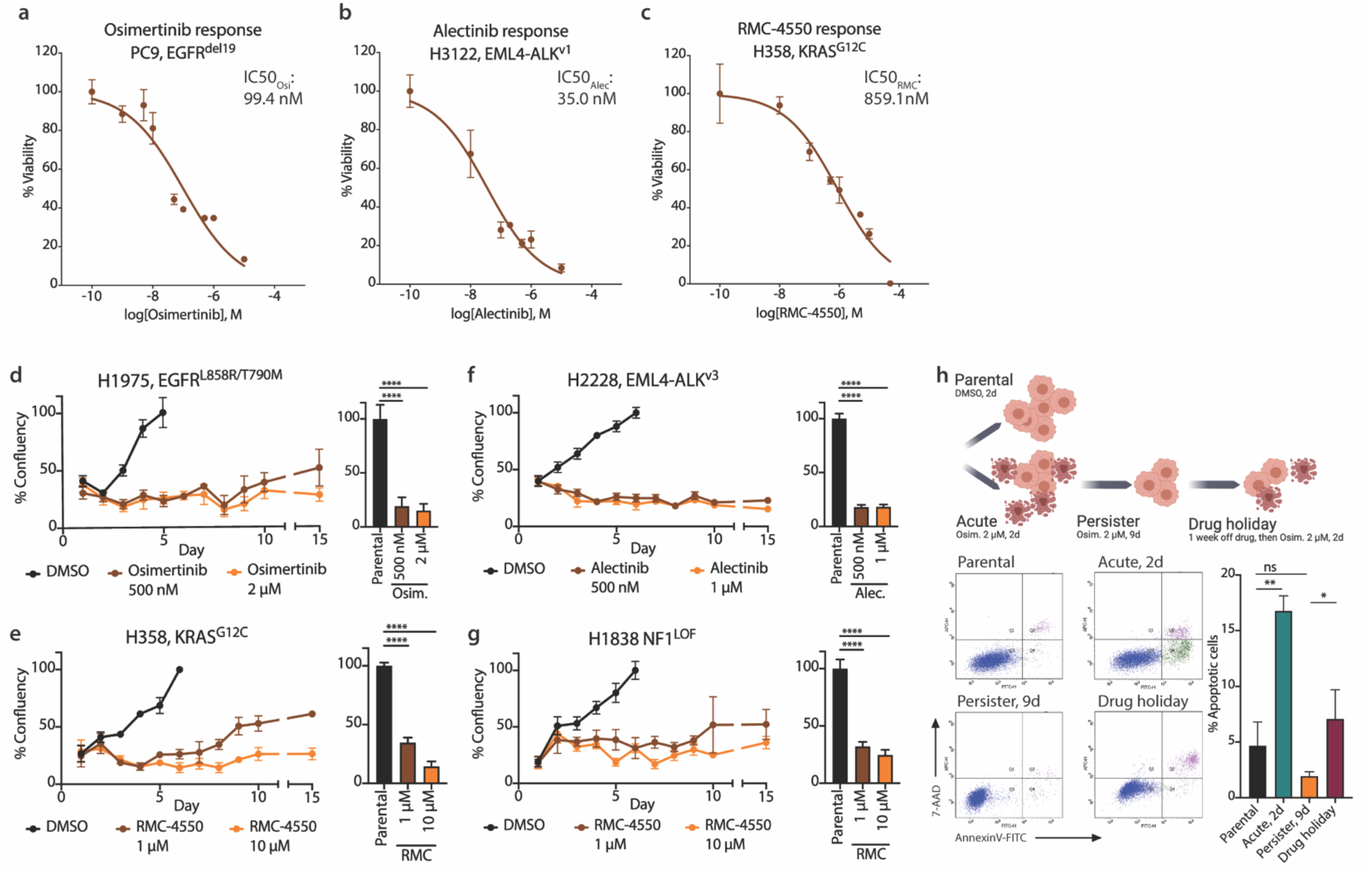
Characterization of drug-tolerant persister cells. (**a**-**c**) Drug response curve to targeted inhibitors across EGFR-mutant PC9 cells treated with osimertinib, ALK fusion-positive H3122 cells treated with alectinib, and KRAS-mutant H358 cells treated with RMC-4550. IC50, half maximal inhibitory concentration. (**d**-**f**) High-content microscopy screen monitoring relative cell numbers in cells treated with targeted inhibitors. Confluency of EGFR-mutant H1975 cells treated with 500 nM (brown) and 2 µM (orange) osimertinib, of ALK-fusion positive H2228 cells treated with 500 nM (brown) and 1 µM (orange) alectinib, of KRAS-mutant H358 cells treated with 1 µM (brown) and 10 µM (orange) RMC-4550 as well as of NF1-mutant H1838 cells treated with 1 µM (brown) and 10 µM (orange) RMC-4550 compared to 0.1 % DMSO control (black), respectively (left). *n* = 6 per datapoint. Comparison of full cell confluency reached in 0.1% DMSO-treated parental cells to persister cell numbers at day 8 of treatment (right). Statistical evaluation by unpaired t-test. **** *p* < 0.0001. (**h**) Monitoring of apoptosis by Annexin-V / 7-AAD staining in EGFR-mutant PC9 cells, including untreated parental cells (0.1 % DMSO), acutely treated cells (2-day osimertinib 2 µM), drug-tolerant persister cells (9-day osimertinib 2 µM), and drug-tolerant persisters that were cultured off-drug for 7 days prior to re-treatment with 2 µM osimertinib for 2 days. *n* = 6 per datapoint. Statistical evaluation by unpaired t-test. Parental vs Acute, ** *p* = 0.0012; parental vs persisters, ns, *p* = 0.0950; parental vs drug holiday, * *p* = 0.0288. Schematic diagram was created with BioRender.com.

**Supplementary Fig. 2.**
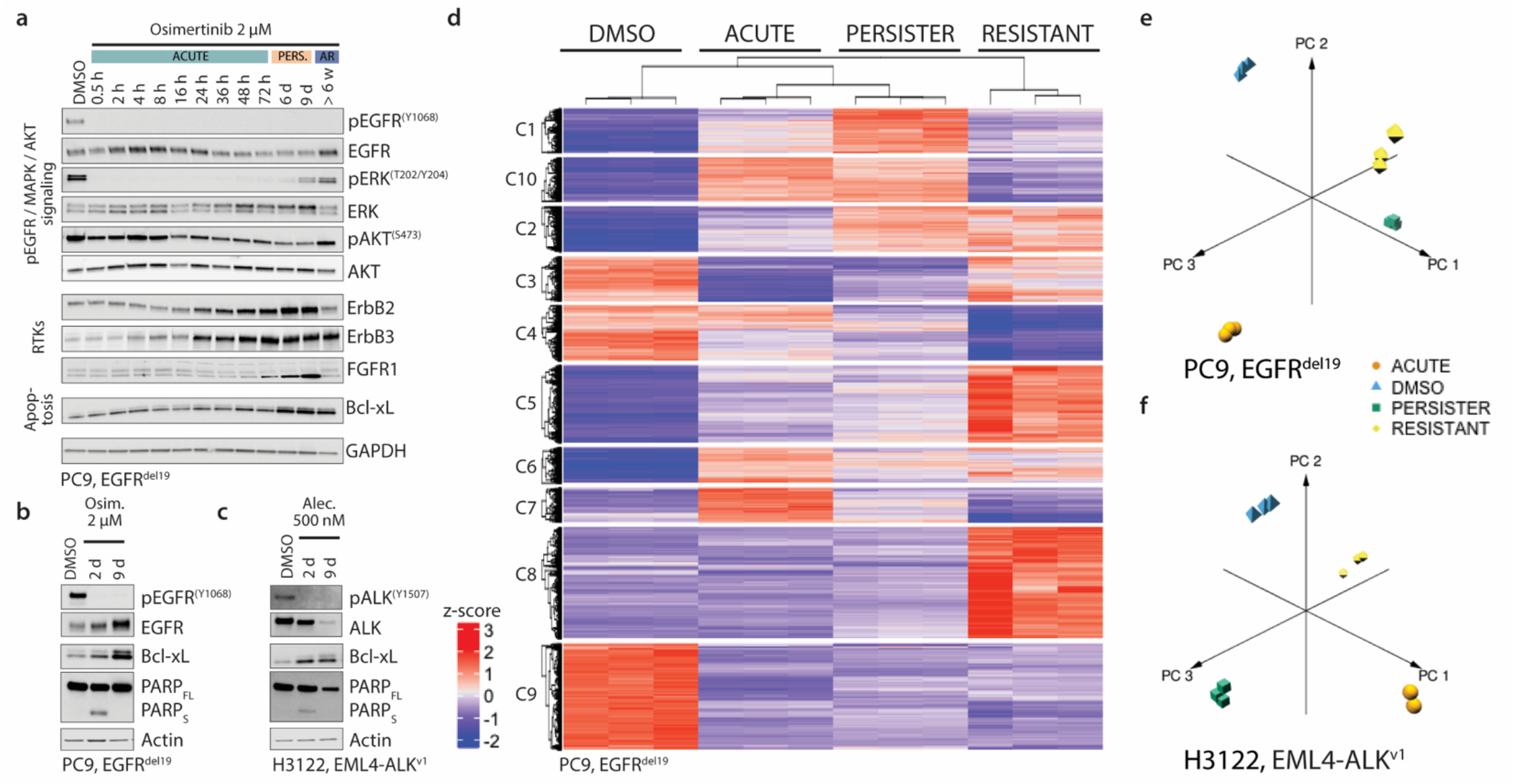
Signaling and transcriptional changes in drug-tolerant persister cells. (**a**) Signaling changes during persister development in EGFR-mutant PC9 cells treated with 2 µM osimertinib, focusing on osimertinib-mediated suppression of EGFR-ERK signaling as well as expression changes for receptor tyrosine kinases ErbB2, ErbB3, FGFR1, and FGFR2 as well as anti-apoptotic Bcl-xL. (**b**-**c**) On-target signaling suppression by targeted inhibitors in acutely treated (day 2) and drug-tolerant persister cells (day 9), monitored for phosphoEGFR-Y1068 in osimertinib-treated PC9 cells and phosphoALK-Y1507 in alectinib-treated H3122 cells. Alterations in apoptosis during persister cell generation with elevated levels of apoptosis-associated cleaved PARP (PARPS) in acutely treated cells (day 2) and absence of cleaved PARP and increase of anti-apoptotic protein Bcl-xL in drug-tolerant persisters (day 9). (**d**-**f**) Changes in RNA expression across 0.1 % DMSO treated parental cells (DMSO), acutely treated cells (ACUTE), drug-tolerant persister cells (PERSISTER), and acquired resistant cells (RESITANT). (**d**) Heatmap analysis of differentially expressed genes in EGFR-mutant PC9 cells and derived acquired resistant PC9-AR cells treated with 2 µM osimertinib. (**e**-**f**) Principal component analysis across treatment states for EGFR-mutant PC9 cells treated with osimertinib and ALK fusion-positive H3122 cells treated with alectinib.

**Supplementary Fig. 3.**
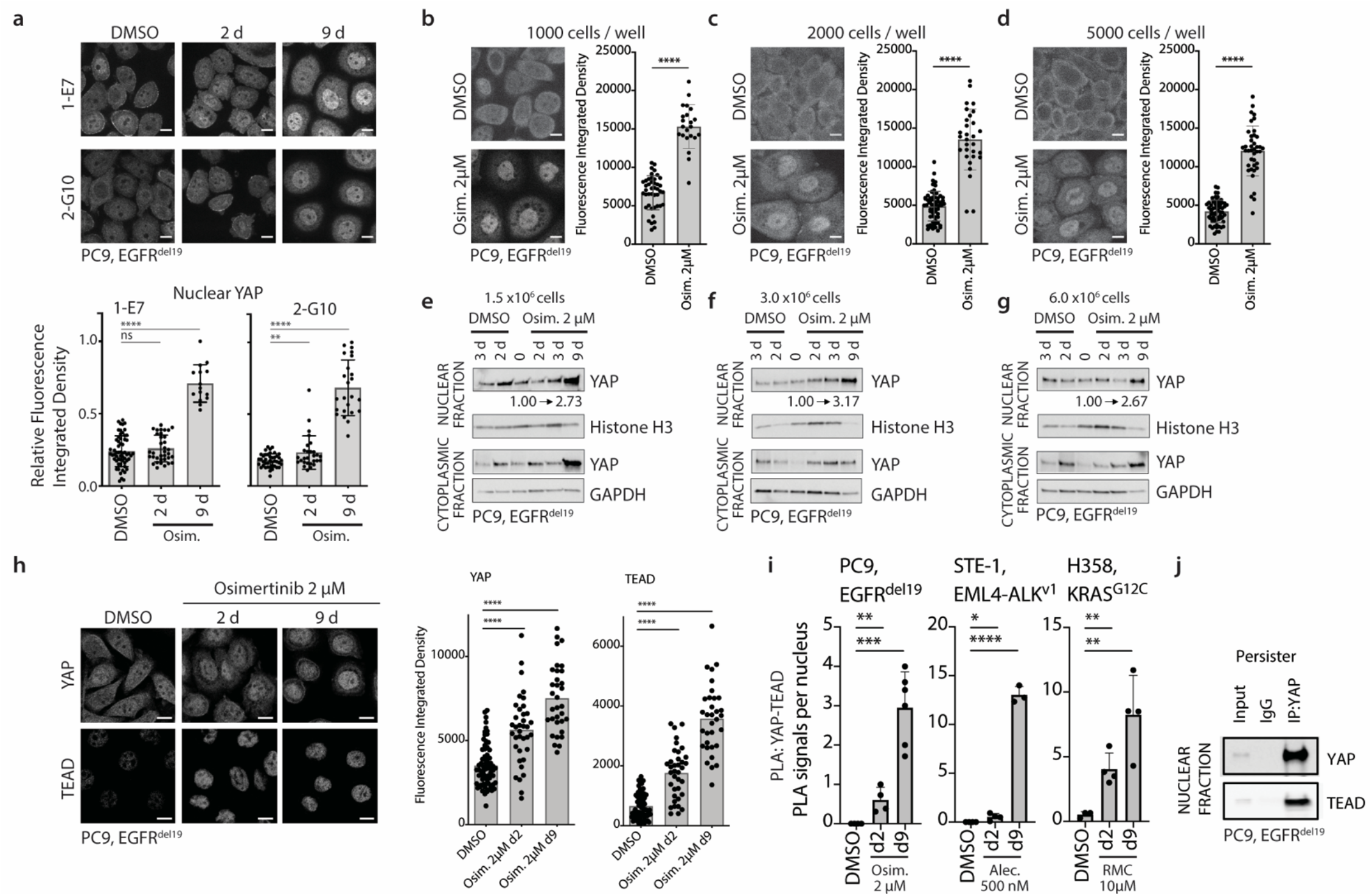
YAP as a functional marker in drug-tolerant persister cells. (**a**) Generation of isogenic, endogenously mNeonGreen (mNG)-tagged YAP PC9 cell lines 1-E7 and 2-G10 and monitoring of changes in YAP nuclear levels by Confocal microscopy. Quantification of fluorescence integrated density per nuclei in 1-E7 and 2-G10 YAP-mNG+ PC9 cells confirms significant increase in YAP nuclear levels in persister cells at day 9. Statistical evaluation by unpaired t-test. 1-E7: DMSO vs 2-day osimertinib (acute), ns, *p* = 0.4058; DMSO vs 9-day osimertinib (persister) **** *p* < 0.0001. 2-G10: DMSO vs 2-day osimertinib (acute), ** *p* = 0.0083; DMSO vs 9-day osimertinib (persister), **** *p* < 0.0001. (**b**-**d**) Analysis of YAP nuclear levels upon osimertinib treatment in PC9 cells seeded at different densities, i.e., sparse with 1000 cells/ well at seeding, intermediate with 2000 cells/ well at seeding, and dense with 5000 cells/ well at seeding in 96-well format, and analyzed at day 5 after seeding by Confocal microscopy. Statistical evaluation by unpaired t-test. **** *p* < 0.0001. (**e**-**g**) Analysis of YAP nuclear and cytoplasmatic levels upon osimertinib treatment in PC9 cells seeded at different densities, i.e., sparse with 1.5 x 10E6 cells/ 150 mm dish at seeding, intermediate with 3.0 x 10E6 cells/ 150 mm dish at seeding, and dense with 6 x 10E6 cells/ 150 mm dish at seeding, and analyzed at indicated timepoints after treatment initiation by immunoblot. (**h**) Nuclear localization of transcriptional co-activator YAP as well as transcription factor TEAD upon treatment of EGFR-mutant PC9 cells with 2 µM osimertinib, analyzed by Confocal microscopy. Representative images (left), scale bar: 10 µm. Quantification of relative integrated density for nuclear levels was performed by automated analysis quantifying the intensity for the protein of interest per nuclei (right). Statistical evaluation by unpaired t-test. **** *p* < 0.0001. (**i**) PanTEAD-YAP proximity ligation assay (PLA) in EGFR-mutant PC9 cells treated with 2 µM osimertinib, ALK fusion-positive STE-1 cells treated with 500 nM alectinib, and KRAS-mutant H358 cells treated with 10 µM RMC-4550. Quantification of PLA signals per nuclei presented as mean value +/- standard deviation, *n* = 3-6 per condition. Statistical evaluation by unpaired t-test with ns, *p* > 0.05; * *p* ≤ 0.05; ** *p* ≤ 0.01; *** *p* ≤ 0.001; **** *p* < 0.0001. (**j**) YAP co-immunoprecipitation (Co-IP) and analysis for concurrent pulldown of TEAD transcription factors in nuclear fractions of 2 µM osimertinib-treated PC9 persister cells at day 9.

**Supplementary Fig. 4.**
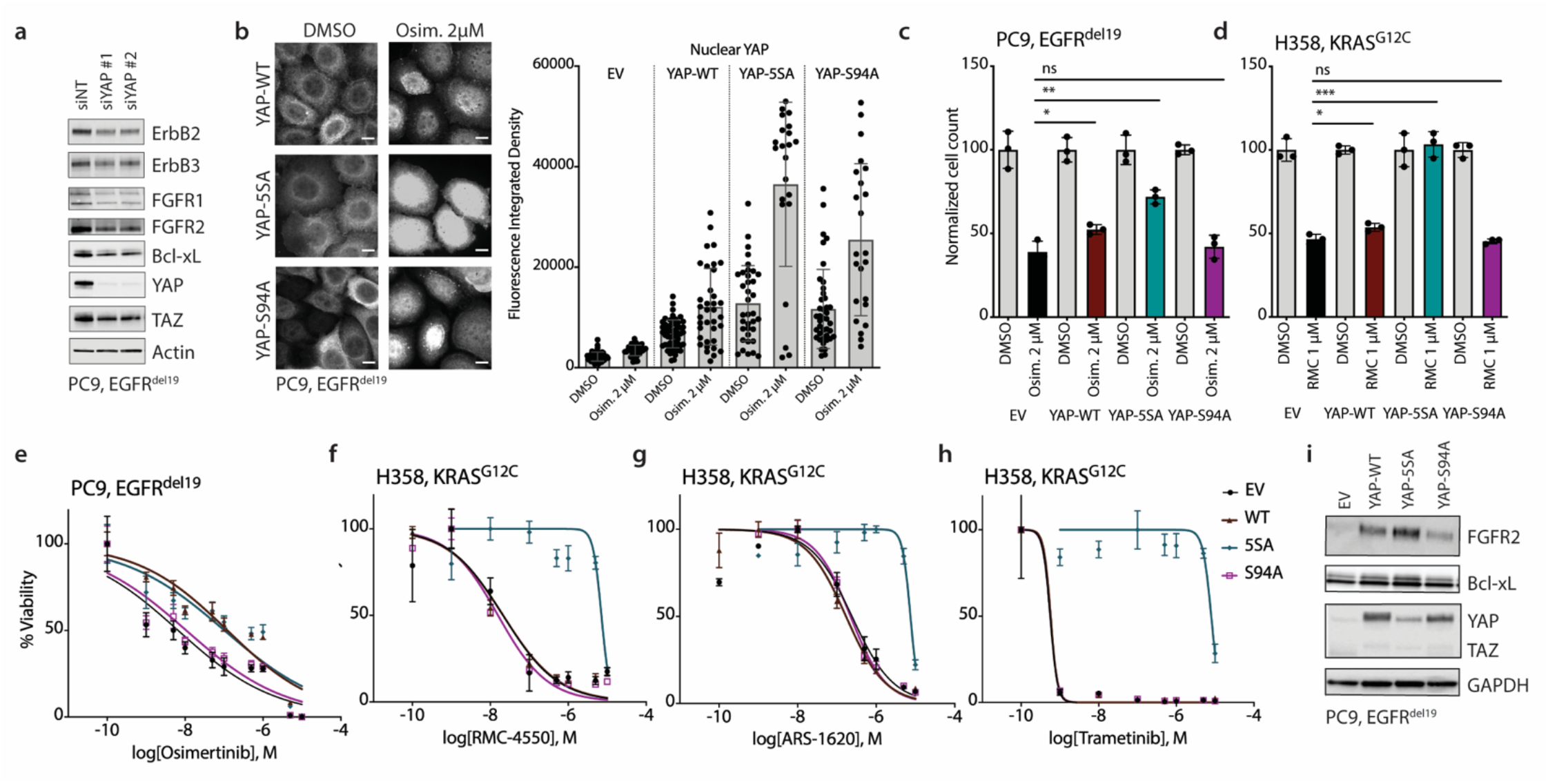
Changes in drug tolerance upon YAP knock-down or overexpression. (**a**) Changes in protein expression of receptor tyrosine kinases ErbB2, ErbB3, FGFR1 and FGFR2 and anti-apoptotic protein Bcl-xL upon siRNA-mediated YAP knock-down in PC9 osimertinib persisters. (**b-d**) YAP nuclear localization (**b**) and relative survival (**c-d**) upon treatment with targeted inhibitors for 48 hours in EGFR-mutant PC9 cells and KRAS-mutant H358 cells overexpressing YAP-WT, hyperactive YAP-5SA, and functionally inactive YAP- S94A, respectively. For (**c-d**), statistical evaluation by unpaired t-test. PC9: empty vector (EV) vs YAP-WT, * *p* = 0.0301; EV vs YAP-5SA, ** *p* = 0.0018; EV vs YAP-S94A, ns, *p* = 0.5990. H358: EV vs YAP-WT, * *p* = 0.0324; EV vs YAP-5SA, *** *p* = 0.0003; EV vs YAP-S94A, ns, *p* = 0.5278. (**e-h**) Treatment response to targeted inhibitors upon overexpression of YAP-WT, hyperactive YAP-5SA, and functionally inactive YAP-S94A, respectively; as shown for the response to osimertinib in EGFR-mutant PC9 cells (**e**) as well as to RMC-4550 (**f**), ARS-1620 (**g**), and Trametinib (**h**) in KRAS-mutant H358 cells. (**g**) Expression changes in YAP target genes Bcl-xL and FGFR2 upon expression of YAP-WT, hyperactive YAP-5SA, and functionally inactive YAP-S94A, respectively, in EGFR-mutant PC9 cells.

**Supplementary Fig. 5.**
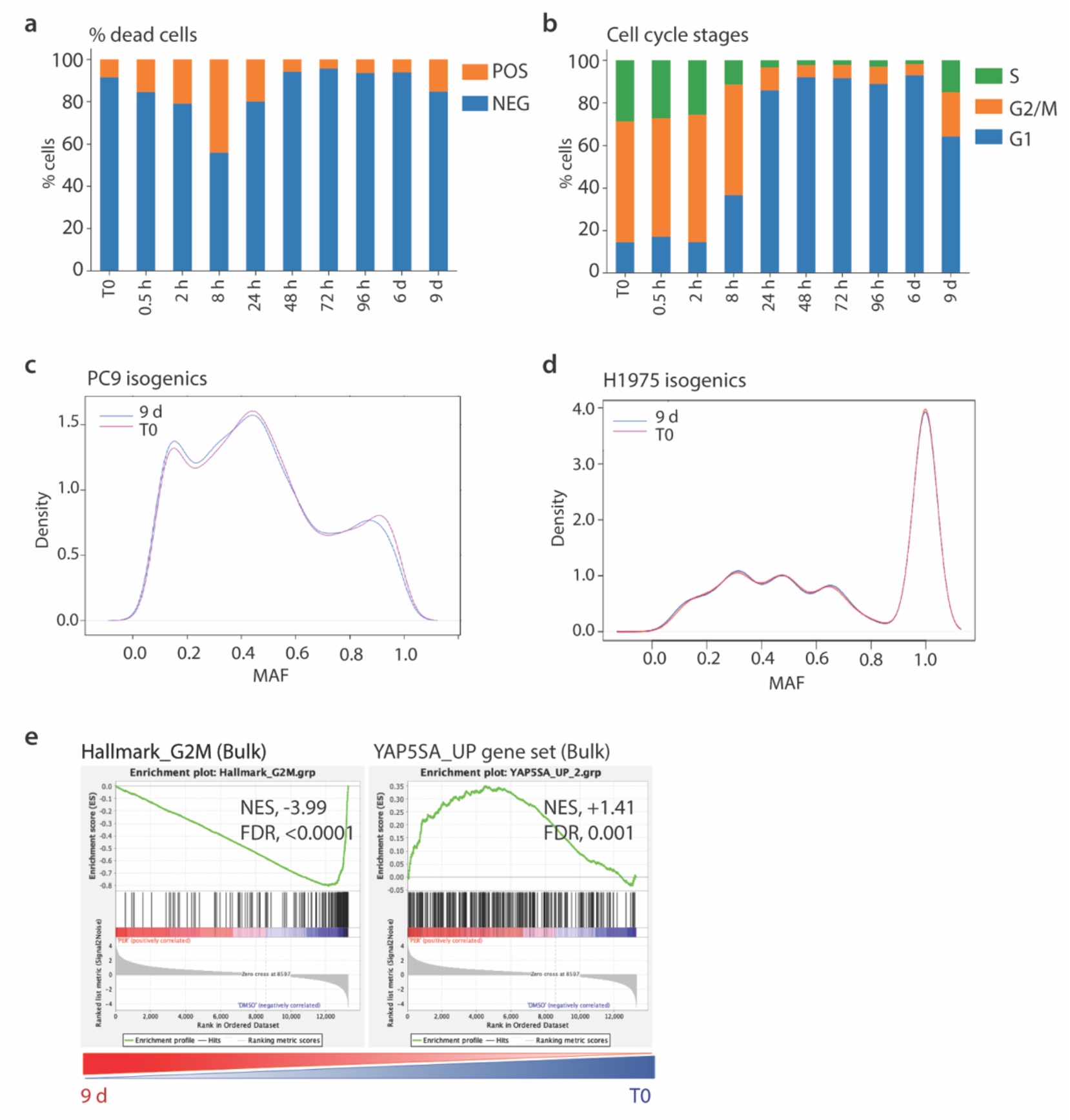
Single cell RNA sequencing trajectory. (**a-b**) Expression changes upon treatment with osimertinib, showing the evaluation of the percentage (%) of dead cells (**a**) and the distribution of cell cycle stages (**b**). (**c**-**d**) Density blot of mutant allele frequencies (MAF) identified by whole exome sequencing and comparing cells prior to treatment (t0) versus at day 9 of osimertinib treatment, showing identical profiles in both conditions and no shift in MAF density under treatment. (**e**) Independent validation of transcriptional changes by bulk RNA sequencing and gene set enrichment analysis for Hallmark_G2M and YAP-5SA_UP gene sets across isogenic EGFR-mutant cell lines, with combined sequencing analysis of isogenic PC9 and H1975 cells (each *n* = 2) and comparing cells prior to treatment (t0) versus at day 9 of osimertinib treatment. *NES*, Nominal Enrichment Score; *FDR*, False Discovery Rate.

**Supplementary Fig. 6.**
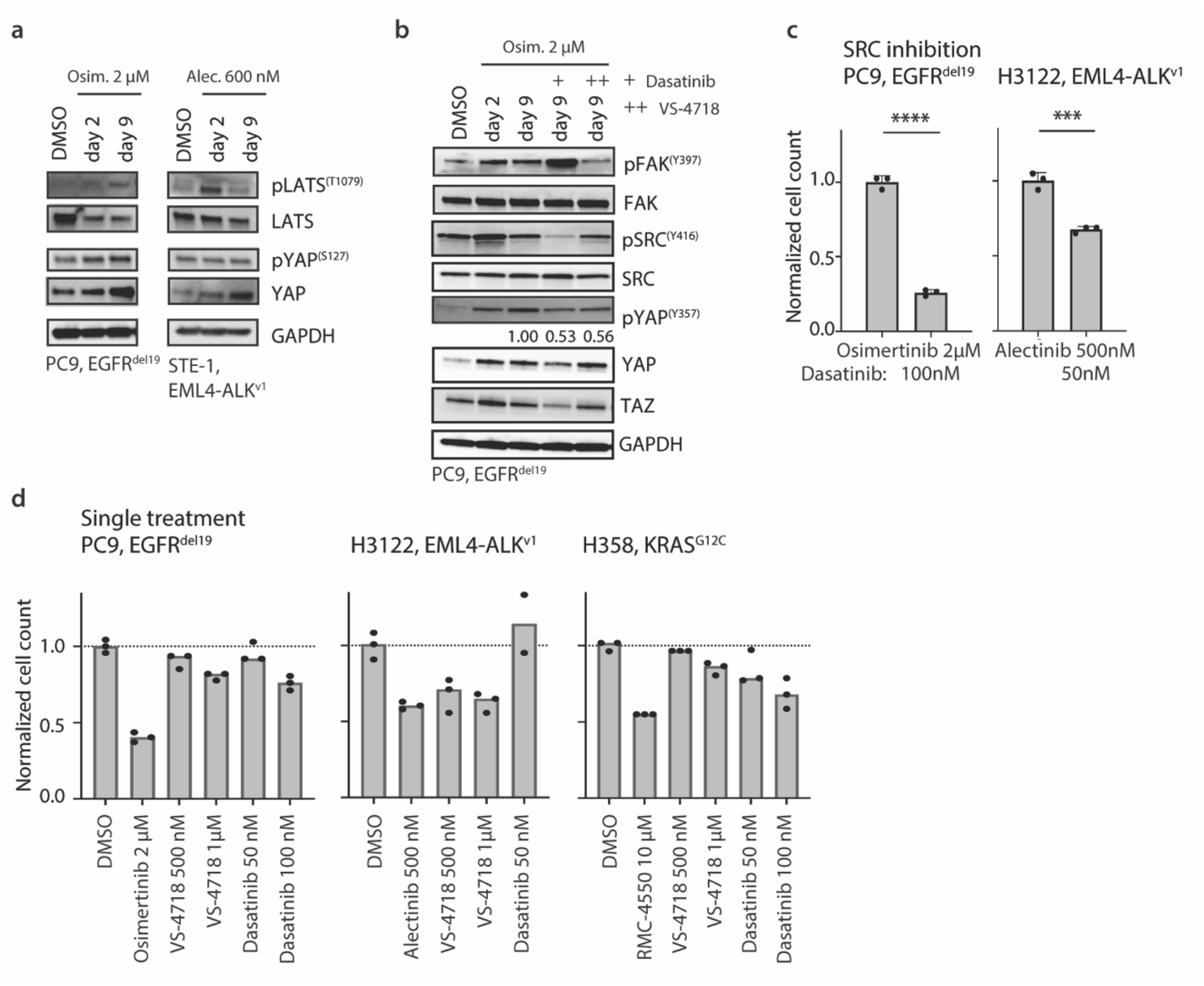
Sensitivity to combinatorial treatment modalities. (**a**) Phosphorylation changes of key Hippo signaling molecules including LATS kinase and the YAP inactivating Ser127 phosphorylation site upon treatment with osimertinib in EGFR-mutant PC9 cells and with alectinib in ALK fusion-positive STE-1 cells. (**b**) Signaling changes during persister development in EGFR-mutant PC9 cells treated with 2 µM osimertinib. Where indicated, short-term treatment of 50 nM Dasatinib (+) or 1 µM VS-4718 (++) has been added for 24 hours prior to harvest. Signaling analysis focuses on combinatorial treatment-mediated suppression of SRC-FAK signaling as well as changes in YAP-Y357 activating phosphorylation. (**c**) Normalized persister cell number for EGFR-mutant PC9 cells upon combinatorial treatment with Src/multikinase inhibitor dasatinib at 50 nM or 100 nM and 2 µM osimertinib during persister cell generation. Statistical evaluation by unpaired t-test. *** *p* = 0.0002; **** *p* < 0.0001. (**d**) Relative cell number compared to 0.1 % DMSO control upon single agent treatment with indicated inhibitors for 48 h.

**Supplementary Fig. 7.**
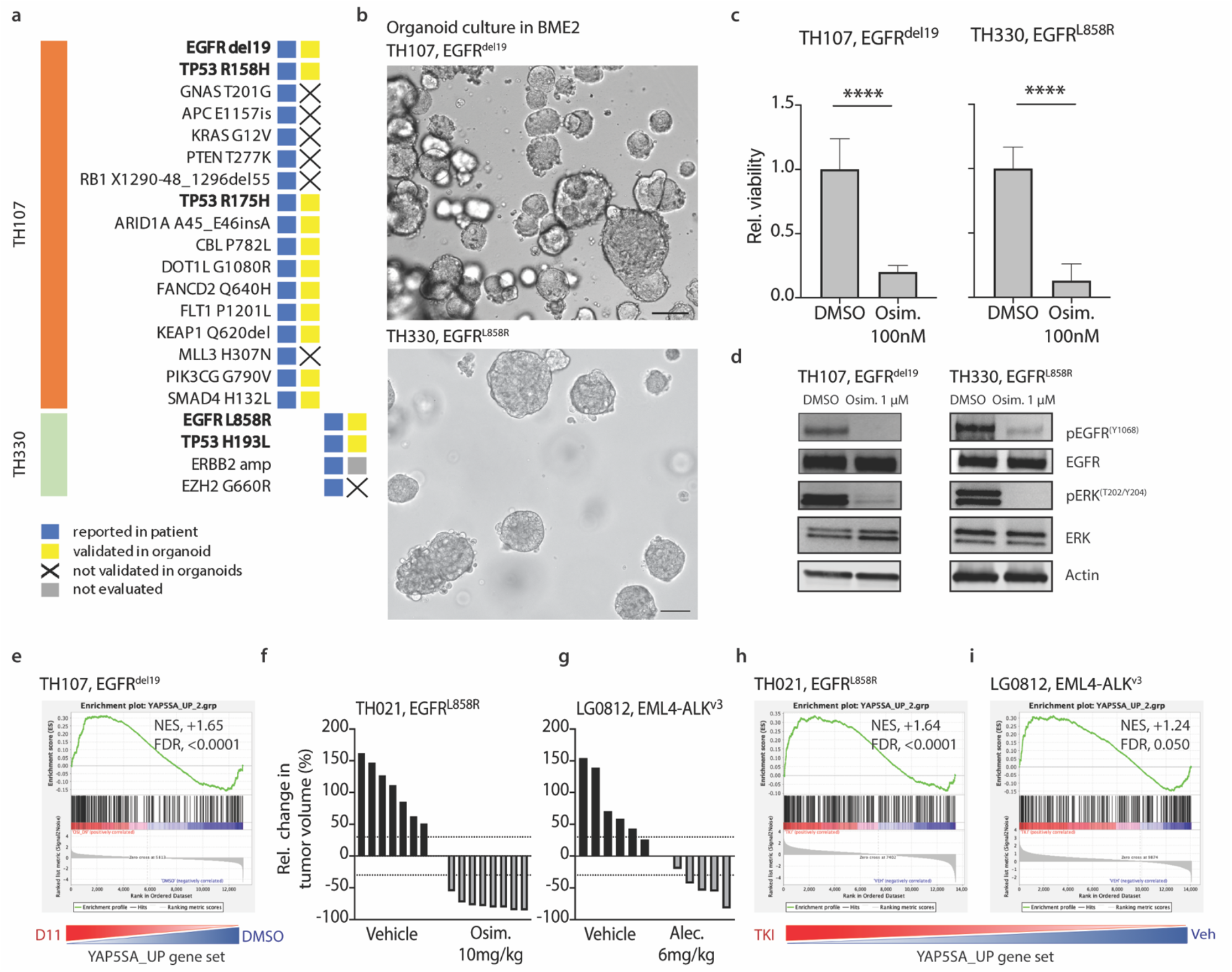
Treatment response in PDO and PDX models. (**a**) Mutation profile of patient specimen and corresponding patient-derived organoid (PDO) models, with indications for the validation status of patient-reported mutation. (**b**) Representative examples of organoid cultures for EGFR-mutant PDOs TH107 (EGFR^del^^19^) and TH330 (EGFR^L858R^); scale bar: 100 µm. (**c**) Sensitivity of EGFR-mutant PDOs to 100 nM osimertinib treatment. (**d**) Suppression of EGFR-ERK signaling upon 1 µM osimertinib treatment in EGFR-mutant PDOs. (**e**) Gene set enrichment analysis for the YAP-5SA_UP gene set using RNAseq expression data of the EGFR-mutant TH107 PDO model, comparing untreated DMSO control (DMSO) versus osimertinib persisters (D11). *NES*, Nominal Enrichment Score; *FDR*, False Discovery Rate. (**f-g**) Relative change in tumor volume for EGFR-mutant PDX TH021 (EGFR^L858R^) treated with 10 mg/kg osimertinib for 7 days (**f**) and for ALK fusion-positive PDX LG0812 (EML4-ALK^v^^3^) treated with 6 mg/kg alectinib for 17 days (**g**), compared to vehicle control. (**h-i**) Gene set enrichment analysis for the YAP-5SA_UP gene set using RNAseq expression data of the EGFR-mutant TH021 PDX model (**g**) and ALK fusion-positive LG0812 PDX model (**h**), comparing vehicle control (VEH) versus treatment group (TKI). *NES*, Nominal Enrichment Score; *FDR*, False Discovery Rate.

**Supplementary Fig. 8.**
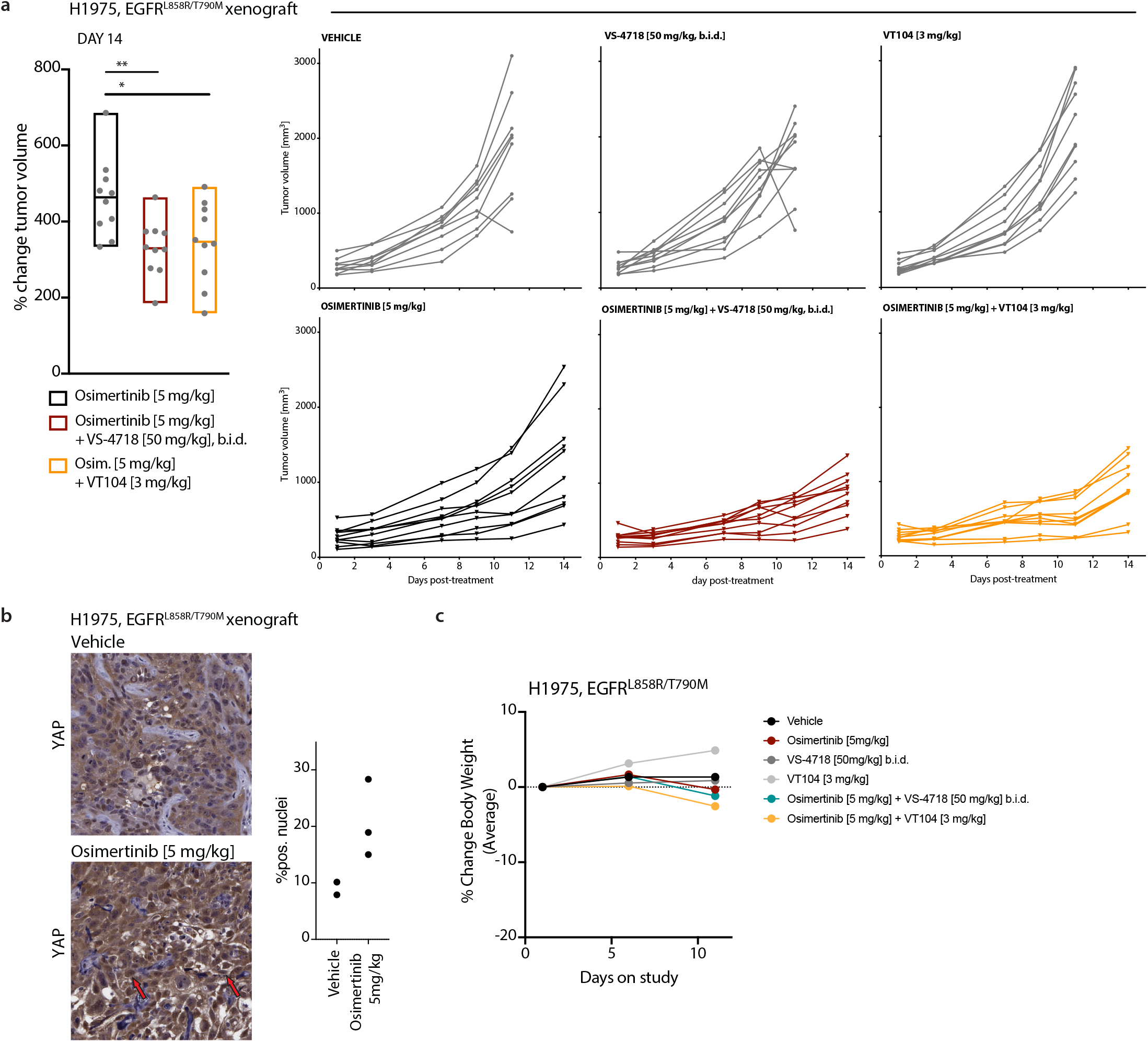
Combinatorial treatment studies in EGFR-mutant H1975 xenograft model. (**a**) Relative tumor volume changes (left) and tumor volume development for individual tumors (right) in an EGFR-mutant H1975 xenograft treatment study across vehicle, 5 mg/kg osimertinib, 50 mg/kg FAK inhibitor VS-4718, 3 mg/kg TEAD inhibitor VT104, and combinatorial 5 mg/kg osimertinib + 3 mg/kg VT104 as well as 5 mg/kg osimertinib + 50 mg/kg FAK inhibitor VS-4718 treatment groups. (**b**) Immunohistochemistry staining for YAP in H1975 xenograft tumor specimens from animals treated with vehicle or osimertinib [5 mg/kg]. Quantification of nuclear levels (% nuclear) by automated image analysis. Arrows indicate cluster of YAP positive tumor cell nuclei. (**c**) Percent change in body weight (BW) for the tumor xenograft treatment study presented in (**a**). Percent change in body weight (BW) was calculated as (BWcurrent - BWinitial)/(BWinitial) x 100%. Data are presented as mean percent body weight change from the day of treatment initiation.

**Supplementary Fig. 9.**
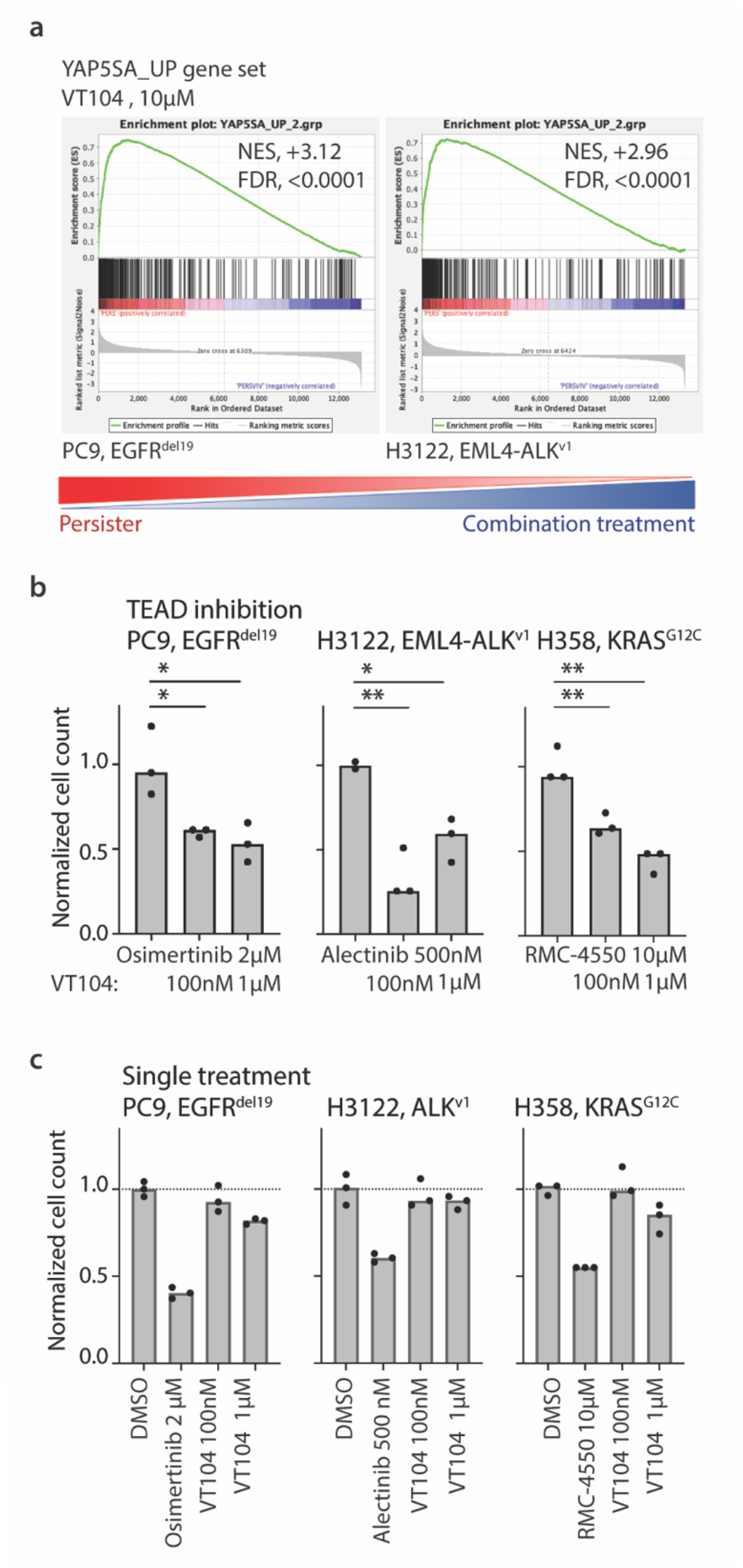
Combinatorial treatment with TEAD inhibitor VT104. (**a**) Gene set enrichment analysis for the YAP-5SA_UP gene set using RNAseq expression data of PC9 osimertinib persister cells and H3122 alectinib persister cells treated with combinatorial 10 µM VT104 for 24 h. *NES*, Nominal Enrichment Score; *FDR*, False Discovery Rate. (**b**) Normalized persister cell number for EGFR-mutant PC9 cells treated with 2 µM osimertinib, ALK fusion-positive H3122 cells treated with 500 nM alectinib and KRAS-mutant H358 cells treated with 10 µM RMC-4550 upon combinatorial treatment with TEAD inhibitor VT104. Statistical evaluation by unpaired t-test with * *p* ≤ 0.05 and ** *p* ≤ 0.01. (**c**) Relative cell number compared to 0.1 % DMSO control upon single agent treatment with indicated inhibitors for 48 h.

**Supplementary Fig. 10.**
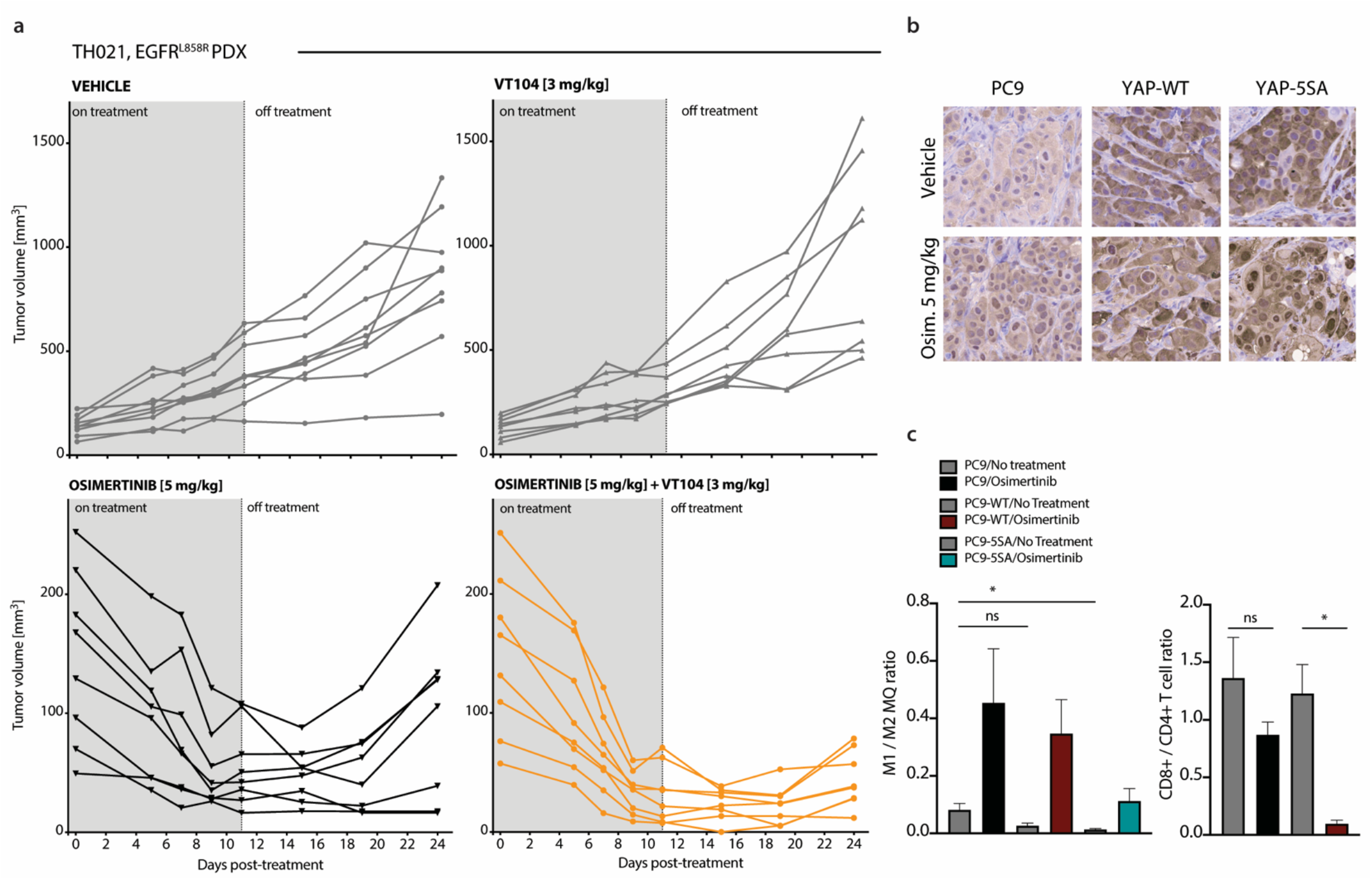
Combinatorial treatment studies in EGFR-mutant TH021 PDX model. (**a**) Tumor volume development for individual tumors in EGFR-mutant TH021 PDX treatment study across vehicle, 5 mg/kg osimertinib, 3 mg/kg TEAD inhibitor VT104, and combinatorial 5 mg/kg osimertinib + 3 mg/kg VT104 treatment groups. (**b**) Immunohistochemistry staining for YAP in humanized PC9 mouse model comparing parental cells, cells expressing YAP-WT and cells expressing hyperactive YAP-5SA. (**c**) Ratios of macrophage (M1 / M2) and T cell (CD4+ / CD8+) populations at treatment endpoint in osimertinib [5 mg/kg] treatment study in humanized PC9 mouse model.

**Supplementary Fig. 11.**
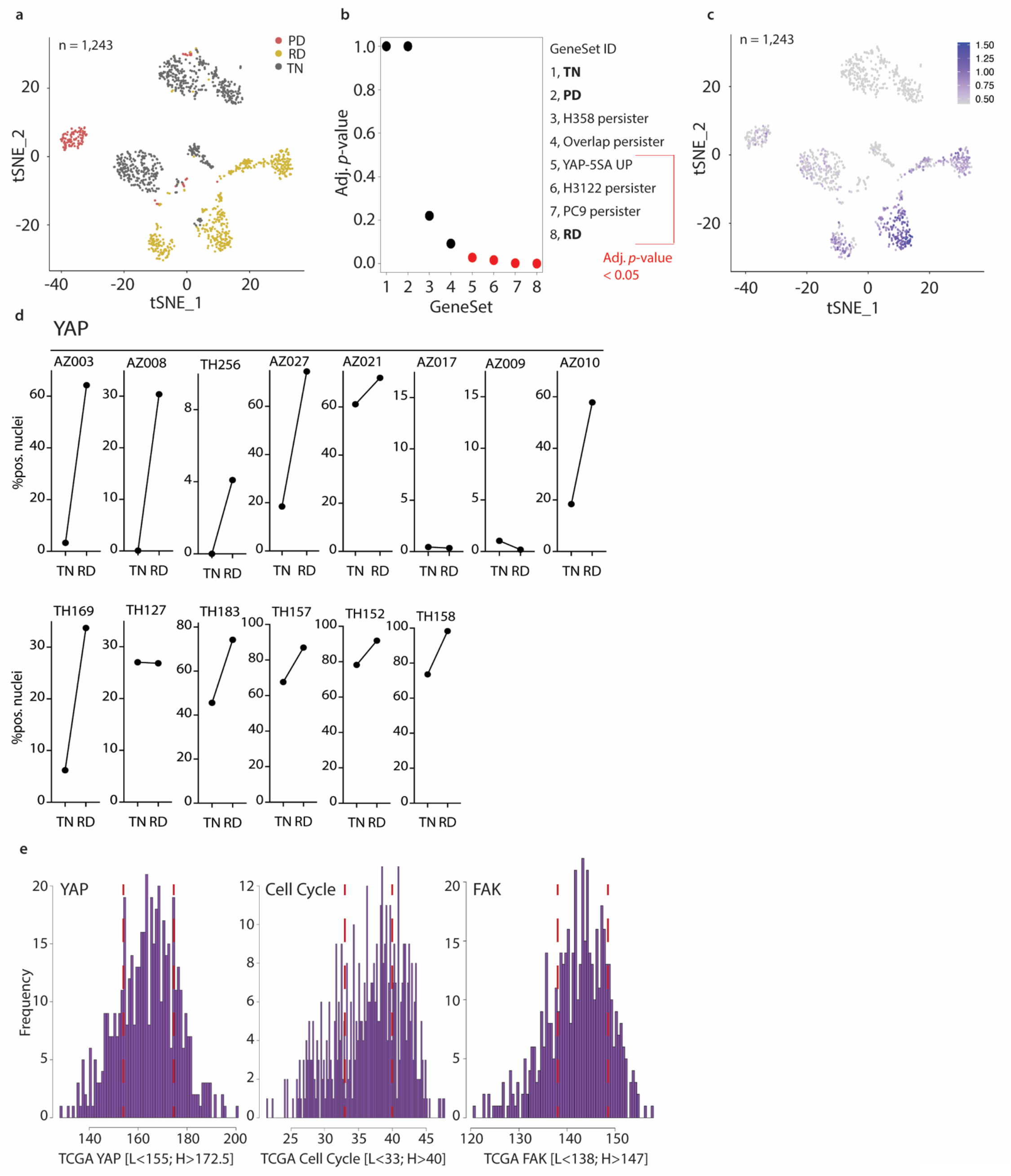
Transcriptional changes and YAP nuclear localization in patient specimens. (**a**) t-stochastic neighbor embedding (t-SNE) plot of all cells derived from primary lung specimen (*n* = 1,243) and colored by treatment timepoint. (**b**) Permutation analysis for indicated gene sets including bulk RNA sequencing data of persister cell line models as well as transcriptional profiles of scRNAseq data from patient specimens collected across TN, RD and PD. Similarity of expression features was determined in relation to scRNAseq profiled at RD. (**c**) Feature plot highlighting the 50% highest expressors for the YAP gene signature across the t-SNE presentation of all cells derived from primary lung specimen (*n* = 1,243). (**d**) Nuclear levels of YAP in matched patient specimens from TN and RD treatment time points. Increase of nuclear YAP levels in 11/14 (∼79 %) matched TN-RD specimens. (**e**) Stratification of expression signature by quartile, with lowest Q1 marking the lowest expression and Q4 the highest expression.

**Supplementary Fig. 12.**
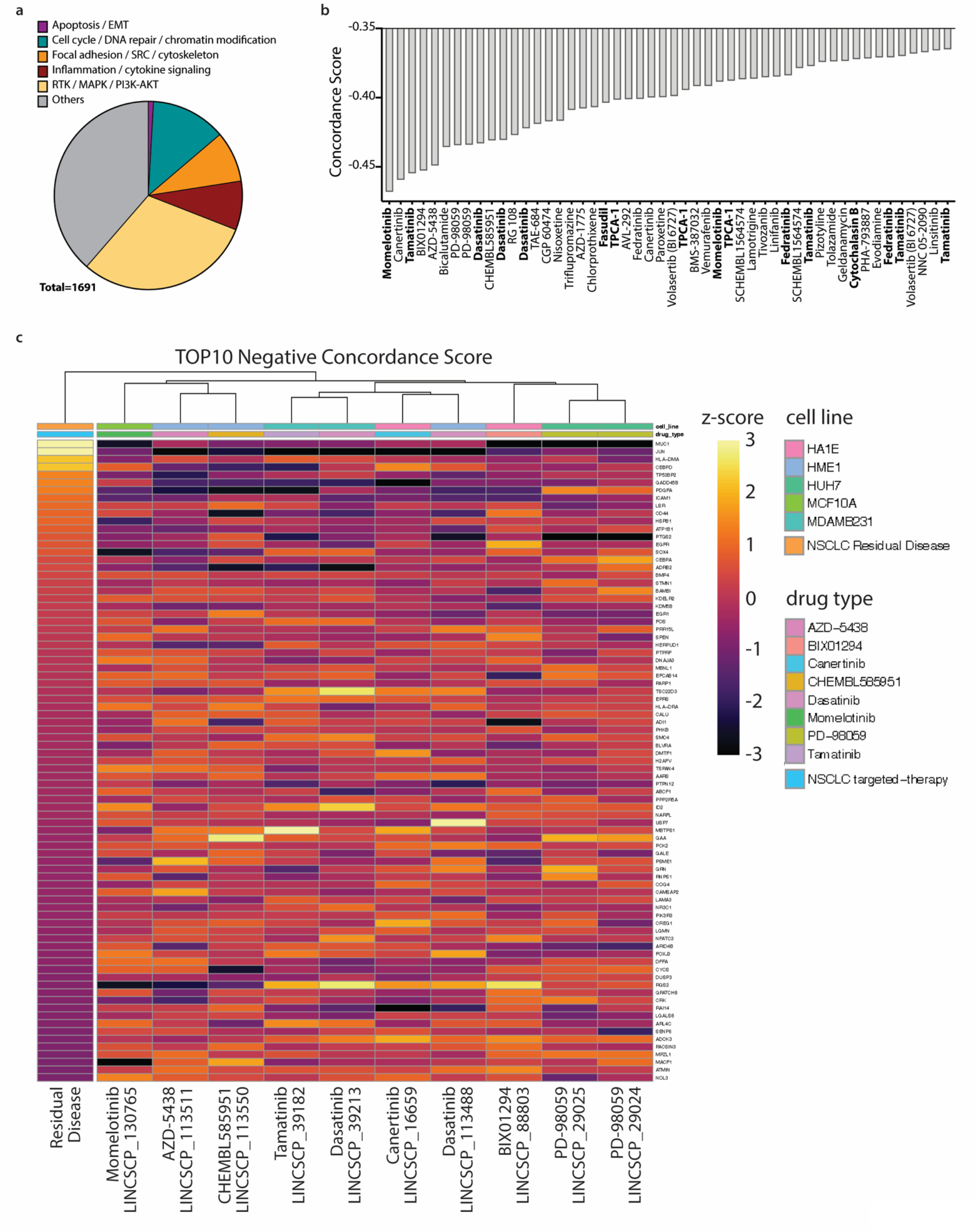
LINCS L1000 concordance analysis. (**a**) Annotation of LINCS L1000 drug / cell line combinations according to molecular processes. Only LINCS L1000 drug / cell line combinations with significant negative concordance score and target gene information are included. (**b**) Bar graph of top 50 annotated LINCS L1000 perturbations with negative concordance scores. Drug names annotated by target gene information within the categories “Focal adhesion / SRC / cytoskeleton” and “Inflammation / cytokine signaling” are highlighted in bold. (**c**) Heatmap presentation of drug-induced expression changes by the top 10 annotated LINCS L1000 perturbations across genes differentially regulated in RD scRNAseq specimens.

